# Functional Annotation of Human Cognitive States using Deep Graph Convolution

**DOI:** 10.1101/2020.04.24.060657

**Authors:** Yu Zhang, Loïc Tetrel, Bertrand Thirion, Pierre Bellec

**Author notes:** **Corresponding Author:** Pierre Bellec, Département de Psychologie, Université de Montréal, 4565, Chemin Queen-Mary, Montréal (Québec) H3W 1W5. **Conflict of interest** The authors declare no competing financial interests.

## Abstract

A key goal in neuroscience is to understand brain mechanisms of cognitive functions. An emerging approach is “brain decoding”, which consists of inferring a set of experimental conditions performed by a participant, using pattern classification of brain activity. Few works so far have attempted to train a brain decoding model that would generalize across many different cognitive tasks drawn from multiple cognitive domains. To tackle this problem, we proposed a ***multidomain*** brain decoder that automatically learns the spatiotemporal dynamics of brain response within a short time window using a deep learning approach. We evaluated the decoding model on a large population of 1200 participants, under 21 different experimental conditions spanning six different cognitive domains, acquired from the Human Connectome Project task-fMRI database. Using a 10s window of fMRI response, the 21 cognitive states were identified with a test accuracy of 90% (chance level 4.8%). Performance remained good when using a 6s window (82%). It was even feasible to decode cognitive states from a single fMRI volume (720ms), with the performance following the shape of the hemodynamic response. Moreover, a saliency map analysis demonstrated that the high decoding performance was driven by the response of biologically meaningful brain regions. Together, we provide an automated tool to annotate human brain activity with fine temporal resolution and fine cognitive granularity. Our model shows potential applications as a reference model for domain adaptation, possibly making contributions in a variety of domains, including neurological and psychiatric disorders.

## Introduction

Identifying brain regions and networks involved in specific cognitive functions has been one of the main goals of neuroscience research (Poldrack, 2006). Modern imaging techniques, such as functional magnetic resonance imaging (fMRI), provide an opportunity to map cognitive function in-vivo. **Traditional fMRI analyses using the general linear model typically implement a direct inference: “if a subject is placed in a certain cognitive state, then a particular pattern of brain regions is activated”. Such findings sometimes get improperly interpreted as a much more powerful statement, called reverse inference: “As soon as a given pattern of brain regions is activated, then participants must have experienced a particular cognitive state”. Poldrack (2011) argues that proper reverse inference can be implemented using a class of techniques called brain decoding, where a given spatial pattern of brain activity is predictive of a given cognitive state, or the stimuli presented to a participant, as was pioneered in the visual domain** (Haxby, 2001). ***In addition, brain decoding should generalize over a large and diverse class of experimental conditions, in order to ensure that the association between a pattern of brain activity and a cognitive state reflects a key property of the corresponding regions, and not a mere reflection of the context where this association was tested*** *(Poldrack, 2011)*. ***The feasibility of such large-scale, multidomain brain decoding is now well established in the literature, e*.*g***. *(Mensch et al., 2017; Poldrack* ***et al***., ***2009; Varoquaux et al***., ***2018), and helps further our understanding of the neural underpinnings of cognitive functions***.

***Previous works on multidomain brain decoding however share a common limitation: decoding is typically based on activation maps, for which task-evoked brain activity is averaged across trials, functional scans or even subjects, in order to generate a single “average” map per condition. So even if a given spatial pattern is found to be predictive of a given cognitive state, that spatial pattern may still be present in specific time points of other conditions. Studies have shown that the temporal dynamics of brain activity may contain discriminative patterns across different cognitive tasks and such brain dynamics are shared among brain regions, or large-scale functional networks*** *(Gonzalez-Castillo et* ***al***., ***2015, 2012; Orban et al***., ***2015). In this work, we aimed to investigate the feasibility of large-scale multi-domain brain decoding over short time series of brain activation***.

Training a brain decoder that distinguishes task conditions across **multiple** cognitive domains **and short time windows** may require the introduction of new machine learning tools, that can handle high-dimensional neural activities distributed across multiple brain systems, and that can at the same time accommodate inter-subject variations in brain organization **and noisy data**. One promising approach is to model the variety of brain dynamics on a brain graph, which provides a network representation of brain organization by associating nodes to brain regions and defining edges via anatomical or functional connections (Bullmore and Sporns, 2009). Based on this architecture, graph signal processing provides a non-linear embedding tool to project brain activities onto Laplacian eigenspaces that integrate spatiotemporal neural dynamics among connected brain regions and networks (Ortega et al., 2018). This approach has been previously used in the neuroscience literature to study the intrinsic organization of brain anatomy and functions. For instance, (Johansen-Berg et al., 2004) separated the human supplementary motor area (SMA) and pre-SMA by mapping the second Laplacian eigenvector of the connectivity matrix derived from diffusion tractography. (Fan et al., 2016) employed a set of Laplacian eigenvectors from the diffusion connectivity profiles and generated the “Brainnectome” whole-brain parcellation Atlas, which consist of 210 cortical and 36 subcortical subregions. Recently, (Margulies et al., 2016) used the graph Laplacian to reveal the gradients of functional organization in the human brain connectome, spanning from primary cortex to the default mode network. In terms of clinical applications, Raj and colleagues found a close correspondence between the Laplacian eigenvectors of whole-brain diffusion tractography profiles generated from healthy subjects and the atrophy patterns measured from Alzheimer’s patients (Raj et al., 2012).

These Laplacian eigenvectors can also be used to build a predictive model of future progression to dementia (Raj et al., 2015). Taken together, these studies suggest great potential of using graph Laplacian in neuroscience research.

In this study, we proposed a multidomain decoding model **over short fMRI time series** by embedding the graph Laplacian with the DNN architecture, called brain graph convolutional networks (GCN). The proposed approach leverages our prior knowledge on brain network organization using graphs, and automatically learns the spatiotemporal dynamics of cognitive processes during model training. Our decoding pipeline (as shown in Fig 1) takes a short series of fMRI volumes as input, maps the fMRI signals onto a predefined brain graph, propagates information of brain dynamics among inter-connected brain regions and networks, generates task-specific representations of recorded brain activities, and then predicts the corresponding task states. We tested the decoding pipeline on the Human Connectome Project (HCP) database by evaluating the performance across 1200 participants and 21 different cognitive tasks at the same time. The performance was compared with a classical brain decoding model, which applies multi-class linear support vector machines on trial-averaged brain activity. Moreover, a valid brain decoding model requires not only a high prediction accuracy but also good interpretability and generalizability. To evaluate whether the decoding inference was based on biologically meaningful **patterns of brain regions**, we generated saliency maps for the input brain response and compared these saliency maps with prior results from the literature on brain anatomy and function. To investigate the temporal sensitivity of the proposed model, we evaluated the performance with time windows of variable length, ranging from a single fMRI volume to the entire block of task trials, and we explored to which extent the performance of the decoding model was constrained by the shape of the hemodynamic response. The stability of the decoding model was finally evaluated by changing the number of subjects used for model training.

**Fig 1.**
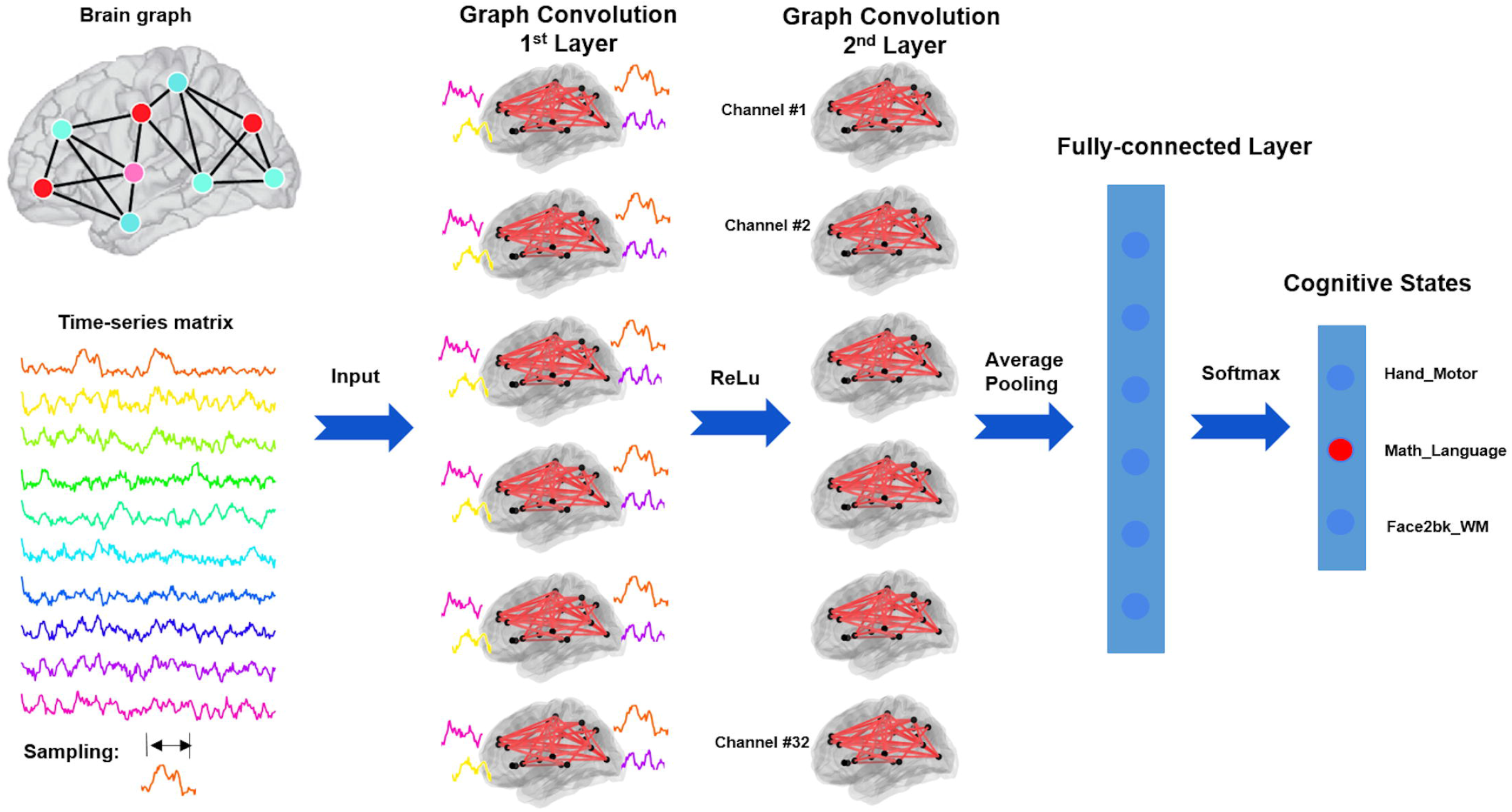
Pipeline of brain state annotation using deep graph convolution network. The proposed state annotation model consists of 6 graph convolutional layers with 32 graph filters at each layer, followed by a global average pooling layer and 2 fully connected layers. The brain graph is constructed by using multimodal cortical parcellation (Glasser et al., 2016) to define the nodes and resting-state functional connectivity to indicate the weights on the edges, both of which were defined based on HCP subjects. A k-nearest-neighbour (k-NN) graph is then built by connecting each brain region only to its 8 neighbors with the highest connectivity. The annotation model takes a short time window of fMRI time series as input, maps the high-dimensional fMRI data onto the brain graph, propagates temporal dynamics of brain response among connected brain regions and networks, generates a high-order graph representation and finally predicts the corresponding cognitive task labels.

## Materials and Methods

### fMRI Datasets and Preprocessing

In this project, we are using the block-design task-fMRI dataset from the Human Connectome Project S1200 release (https://db.humanconnectome.org/data/projects/HCP_1200). The minimal preprocessed fMRI data of the CIFTI format were used, which maps individual fMRI time-series onto the standard surface template with 32k vertices per hemisphere. The preprocessing pipelines includes two steps (Glasser et al., 2013): 1) fMRIVolume pipeline generates “minimally preprocessed” 4D time-series that includes gradient unwarping, motion correction, fieldmap-based EPI distortion correction, brain-boundary-based registration of EPI to structural T1-weighted scan, non-linear (FNIRT) registration into MNI152 space, and grand-mean intensity normalization. 2) fMRISurface pipeline projects fMRI data from the cortical gray matter ribbon onto the individual brain surface and then onto template surface meshes, followed by surface-based smoothing using a geodesic Gaussian algorithm. Further details on fMRI data acquisition, task design and preprocessing can be found in (Barch et al., 2013; Glasser et al., 2013).

The task fMRI data includes seven cognitive tasks, which are emotion, gambling, language, motor, relational, social, and working memory. In total, there are 23 different experimental conditions. Considering the short event design nature of the gambling trials (1.5s for button press, 1s for feedback and 1s for ITI), we evaluated the decoding models (see the pipeline section below) with and without the two gambling conditions and found a much lower precision and recall scores for gambling task (average f1-score = 61%) than other cognitive domains (average f1-score > 91%). In the following experiments, we excluded the two gambling conditions and only reported results on the remaining 21 cognitive states. The detailed description of the tasks can be found in (Barch et al., 2013). A summary table is also shown in Table 1.

**Table 1.** Scanning parameters and experimental designs of HCP task-fMRI dataset.

| **Task Domains** | **#Subjects** | **#Runs** | **#Volumes per run** | **#Trials per run** | **#Conditions** | **Minimal duration per block (sec)** |
| --- | --- | --- | --- | --- | --- | --- |
| Working memory | 1085 | 2 | 405 | 8 | 8 | 25 |
| Motor | 1083 | 2 | 284 | 10 | 5 | 12 |
| Language | 1051 | 2 | 316 | 8 | 2 | 10 |
| Social Cognition | 1051 | 2 | 274 | 5 | 2 | 23 |
| Relational processing | 1043 | 2 | 232 | 6 | 2 | 16 |
| Emotion | 1047 | 2 | 176 | 6 | 2 | 18 |

### Baseline decoding methods

We applied multi-class linear support vector machines (SVC) and random forests (RF) as our baseline methods for brain decoding, which were the most frequently used classification methods in previous brain decoding studies (Li and Fan, 2019; Varoquaux et al., 2018). ***For SVC, we evaluated both linear and nonlinear (RBF) kernels, using the implementations from sklearn. We chose the one-vs-rest (‘ovr’) decision function to handle multi-classes and used a grid search strategy to locate the optimal C parameter: [0*.*001, 0*.*01, 0*.*1, 1***., ***10***., ***100*.*] For RF, we evaluated different settings of the classifier including depth of trees: [4***,***16***,***64***,***256***,***1024] and number of trees: [100***,***2000]. The baseline decoding models took the averaged BOLD signals from every 10s of task blocks as inputs***. We only reported the best decoding performance for each baseline method after the hyperparameter search.

### Convolutional Neural Networks on Brain Graphs

Graph Laplacian and graph signal processing (GSP) provides a generalized framework to analyze data defined on irregular domains, for instance social networks, biological interactions and brain graphs. A brain graph captures a network representation of brain organization by associating nodes with brain regions and defining edges via anatomical or functional connections (Bullmore and Sporns, 2009). Based on this representation, a non-linear embedding tool can be used to project brain activity from large-scale noisy measures in the spatial domain to low-dimensional representations in the spectrum domain (Ortega et al., 2018). This method has gained more and more attention in neuroscience studies, for instance parcellating brain areas (Fan et al., 2016; Johansen-Berg et al., 2004), identifying functional areas and networks (Atasoy et al., 2016; Craddock et al., 2012), and generating connectivity gradients (Margulies et al., 2016). Recently, studies have found that, by decomposing the task-evoked fMRI signals using GSP, the resultant graph representations strongly associated with cognitive performance and learning (Huang et al., 2016; Medaglia et al., 2018). These findings brought new opportunities for the application of GSP on neuroimaging analysis.

### Definition of Brain graph

Starting with assigning a brain signal *x* ∈ ℝ^*N* ×*T*^, i.e. a short time-series with duration of, *T* to each of *N* brain regions, GSP maps the recorded brain activity onto a weighted graph 𝒢 = (𝒱, *ε*, *W*) that defines the network architecture among a set of brain regions. The set 𝒱 is a parcellation of cerebral cortex into *N* regions, and *ε* is a set of connections between each pair of brain regions, with its weights defined as *W*_*i,j*_. Many alternative approaches can be used to build such brain graph 𝒢, for instance using different brain parcellation schemes and constructing various types of brain connectomes (for a review, see (Bullmore and Sporns, 2009)). Here we used the multimodal cortical parcellation defined based on 210 subjects from Human Connectome Project (HCP) (Glasser et al., 2016), which delineates 180 areas per hemisphere, bounded by sharp changes in cortical architecture, function, connectivity, and topography. The edges between brain areas were estimated by calculating the group averaged resting-state functional connectivity (RSFC) based on minimal preprocessed resting-state fMRI data from *N*=1080 HCP subjects (Glasser et al., 2013). Additional preprocessing steps were applied before the calculation of RSFC, including regressing out the signals from white matter and csf, and bandpass temporal filtering on frequencies between 0.01 to 0.1 HZ. Functional connectivity was first calculated on individual brains using Pearson correlation and then normalized using Fisher z-transform before averaging among the entire group of subjects. After that, a k-nearest-neighbour (k-NN) graph was built by only connecting each brain region to its 8 neighbours with highest connectivity (see Fig 1-Supplement 1 as an example).

### Graph Laplacian and Graph Fourier transform

The spectral analysis of the graph signals relies on the graph Laplacian, which maps the signal distributions from the spatial domain to the graph spectral domain and decomposes the signals into a series of graph modes with different frequencies. Specifically, the normalized graph Laplacian matrix is defined as:

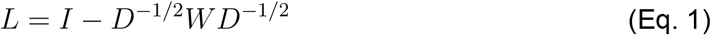

where *D* is a diagonal matrix of node degrees and *I* is the identity matrix. As we assume the weights to be undirected and symmetric, the matrix *L* can be factored as, *U* Δ *U* ^*T*^ where *U* = (*u*_0,_…,*u*_*N−* 1_ is the matrix of Laplacian eigenvectors and is also called graph Fourier modes, and Δ = diag(λ_0_, …, λ_*N*−1_) is a diagonal matrix of the corresponding eigenvalues, specifying the frequency of the graph modes. In other words, the eigenvalues quantify the smoothness of signal changes on the graph, while the eigenvectors indicate the patterns of signal distribution on the graph. This eigendecomposition can be interpreted as a generalization of the standard Fourier basis onto a non-Euclidean domain (Bronstein et al., 2017; Shuman et al., 2013). Based on the eigendecomposition, the graph Fourier transform is defined as 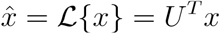 and its inverse as 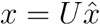, where *x* ∈ ℝ^*N* × *T*^ is the graph signal and 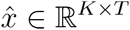 is the transformed signal with *K* selected eigenvectors or graph modes. The Laplacian matrix and these transformations are the fundamental basis of graph signal processing and graph convolutional networks.

### Graph Convolutional Networks: spectral

Recently, graph convolutional neural networks (GCN) was proposed to merge graph signal processing with the deep neural network architecture (Bruna et al., 2013). The key step is to generalize the convolution operations onto the graph domain. Instead of calculating a weighted sum among the spatial neighbours in the Euclidean space as in a classical convolutional neural network (Krizhevsky et al., 2012), GCN generates a linear combination of graph Fourier modes across different frequencies by using graph filters. Specifically, the convolution between the graph signal *x* ∈ ℝ^*N* × *T*^ and a graph filter *g_θ_* ∈ ℝ^*N* × *T*^ (independent weight matrix for each temporal channel) based on graph 𝒢, is defined as their element-wise Hadamard product in the spectral domain, i.e.:

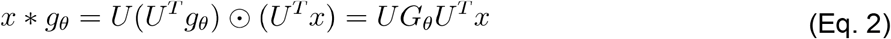

where *G*_θ_ = diag (*U*^*T*^*gθ*) and *θ* indicates a parametric model for *gθ*, and *U*^*T*^*x* is actually projecting the graph signal onto the full spectrum of graph modes. With different choices of *θ* GCN learns different types of graph filters and finds the optimal graph representations of the input signals for a given task.

### Graph Convolutional Networks: ChebNet

To avoid calculating the spectral decomposition of the graph Laplacian, especially for large-scale graphs, ChebNet convolution (Defferrard et al., 2016) uses a truncated expansion of the Chebychev polynomials, which are defined recursively by:

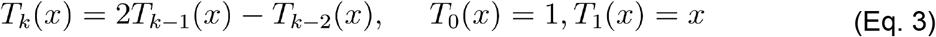

Consequently, the graph convolution is defined as:

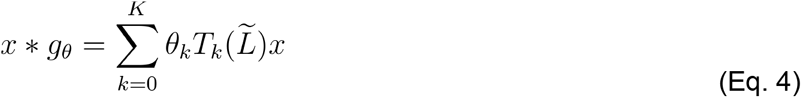

where 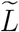 is a normalized version of graph Laplacian, equals to 2*L*/λ_max_ − *I*, with λ_max_ being the largest eigenvalue, *θ*_*k*_ is the parameter to be learned for each order of the Chebychev polynomials.

### Graph Convolutional Networks: 1st-order

Kipf and colleagues (Kipf and Welling, 2016) introduced a simplified version of GCN by taking a first-order approximation of the above Chebychev polynomial expansion and: λ_max_ ≈ 2;

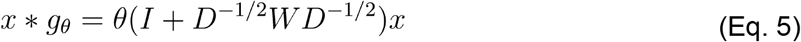

where *θ* is ***a parameter vector with one parameter to be learned at each time point*** and *W* is the weight matrix for brain connectome.

### Graph Convolutional Networks: multi-layer

Complex signal representations can be learned by stacking multiple layers of graph convolutions. The output of a graph convolution layer is defined as:

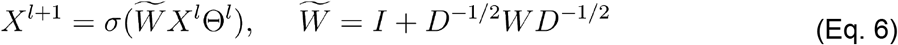

***where*** *X*^*l*^∈ ℝ^*N* × *C;*^ ***denotes the input graph signals on layer*** *l*, ***with*** *N* ***brain regions and*** *C* ***input channels***, *X*^*l*+1^∈ ℝ^*N* ×*F*^ ***denotes the output of layer*** *l* ***with*** *N* ***brain regions and*** *F* ***graph filters. Θ***^*l*^∈ ℝ^*C*×*F*^ ***is the parameters of graph convolution kernel on layer*** *l* ***with*** *C* ***input channels and outcome filters. These parameters are shared among all brain regions. To be noted that in the first graph convolution layer, the input channels*** *C* ***is equal to the time window of input fMRI time-series***. denotes an activation function, such as the ReLU (*x*) = max(0,*x*). It’s worth noting that the first-order GCN only takes into account the direct neighbours for each brain region which are indicated by the adjacency matrix of the graph. By stacking multiple GCN layers, we could propagate brain activity among the *k*_*th*_ -order neighbourhood, i.e. connecting two nodes by passing *k* − 1 other neighbours in between, with *k* is the number of convolution layers. In the following analysis, we are using the multi-layer architecture of 1st-order GCN for brain decoding. Here, ***we used 6 GCN layers according to the 6 degrees of separation in the small world theory such that the model integrates brain dynamics of any two brain regions in a maximum of six steps***.

### Brain State Annotation pipeline

We propose a brain state annotation model consisting of 6 graph convolutional layers with 32 graph filters at each layer, followed by a global average pooling layer and 2 fully connected layers (256, 128 units). Specifically, the input to the first GCN layer is a short series of fMRI volumes by treating multiple time steps as input channels (*C* = *T*), i.e. *X* ^1^ ∈ ℝ^*N*×*T*^ being a 2D matrix *N* consisting of brain regions and *T* time steps. During model training, the first GCN layer learns various versions of the spatiotemporal convolution kernel (integrating information from graph neighbors in space, and training separate kernels for each time step) for fMRI time-series, as a replacement of the canonical hemodynamic response function (HRF). ***Examples of temporal filters learned in the first GCN layer was shown in Fig 7-Supplement 2***. The model takes a short series of fMRI data as input, propagates information among inter-connected brain regions and networks, generates a high-order graph representation and finally predicts the corresponding cognitive labels as a multi-class classification problem. An overview of the fMRI decoding model was illustrated in Fig 1.

The entire dataset was split into training (70%), validation (10%), test (20%) sets using a subject-specific split scheme, which ensures that all fMRI data from the same subject was assigned to one of the three sets. Specifically, for each subject and each cognitive domain, individual fMRI time-series on the 64k surface template (including both hemispheres) was first mapped onto the 360 areas of Glasser atlas (Glasser et al., 2016), by averaging the BOLD signals within each parcel. The time-series of each task trial was extracted and saved into a 2D matrix, by first realigning fMRI signals with experimental designs of event tasks using task onsets and durations and then cutting the time-series into bins of selected time window (see *Time window* section below). Next, the time-series matrices from all training subjects were collected into a pool of data samples. At each step of model training, a set of data samples (e.g. 128 time-series matrices) was input to the decoding model and the parameter matrix Θ^*l*^ of each layer were optimized through gradient descent. After all data samples have been trained (i.e. finishing one epoch), the model was then evaluated on the samples from the validation set before the next epoch started. The best model with the highest prediction score on the validation set was saved and then evaluated separately on the test set. There are mainly two types of decoding models used in this study, either training by exclusively using fMRI data from a single cognitive domain or combining fMRI data from multiple cognitive domains. The rectified linear unit (ReLU) function (Maas et al., 2013) was used as the activation function for all layers except the last layer where the softmax function was used to predict the cognitive labels. The network was trained for 100 epochs with the batch size set to 128. We used Adam as the optimizer with the initial learning rate as 0.001. Additional l2 regularization of 0.0005 on weights was used to control model overfitting and the noise effect of fMRI signals. Dropout of 0.5 was additionally applied to the neurons in the last two fully connected layers. The implementation of the GCN model was based on Tensorflow 1.12.0, and was made publicly available in the following repository: https://github.com/zhangyu2ustc/GCN_fmri_decoding.git.

### Time window of fMRI data

As mentioned above, we treated the fMRI time windows as multiple input channels in the first layer of the GCN model. There are several benefits of using multiple input channels. First, the network is enriched with more low-level graph filters, which provides more diverse features for the high-level graph convolutions. Second, with long enough fMRI time series, the network trains its own versions of the convolution kernel based on the fluctuation of task-evoked BOLD signals, as a replacement of the canonical HRF, that typically includes a small initial dip, followed by a dominant peak at 4-6s after the onset of neural activity, and then a variable post-stimulus undershoot around 8-12s after onset (Buxton et al., 1998). In the meantime, the different shapes of fluctuations are also informative regarding the cognitive states and could help the GCN model in state annotation.

To test this effect, we first trained a GCN model with only one input channel, i.e. using a single fMRI volume as input and predicting the cognitive label associated with that fMRI volume. It’s worth noting that, according to this design, each fMRI volume during the task event (from task onset to the end of each task trial) was treated as an independent data sample. As a result, brain response at different stages of task-evoked hemodynamic response was embedded by learning multiple graph filters during model training. Thus, we could evaluate the performance of GCN annotation as a function of time-elapsed-from-onset, ranging from 0 to the length of the entire task trial. F1-score (Powers, 2011) was used as a measure of the prediction accuracy, which is the harmonic average of the precision and recall, with its best value at 1 (perfect precision and recall) and worst at 0.

Considering the low temporal signal-to-noise ratio of fMRI acquisition, especially for a single fMRI volume, we tested the same procedure with 6s of fMRI time series which includes 8 input channels at the first convolution layer. Specifically, the fMRI time-series of all task trials were first cut into non-overlapping mini-blocks of 6s time window. For instance, as for the 12s movement trials from the motor task, we compared the GCN performance in predicting different types of movements at time bins of 0-6s vs 6-12s after task onset. These short bins of time-series were treated as independent data samples during model training. For those task trials shorter than 12s, we applied a neighborhood wrapping method by using numpy.take. For instance, some of the mathematical task trials only last for 10s. In order to match the time window of the input fMRI data, we repeated the fMRI scan at the end of the task trial several times matching for 12s.

Other time windows were also evaluated, ranging from a single fMRI volume (0.72s) to the minimal duration of all task trials (10s) at a step of two TRs (1.4s). The decoding accuracies on the test set were fitted with an exponential function and summarized by averaging the performance within each cognitive domain.

### Size of the dataset

The Human Connectome Project recruits 1200 healthy participants. It also provides us an opportunity to evaluate the sample size effect, i.e. how many independent subjects were sufficient to reach the stable performance of GCN. To test that, we scanned over the entire task-fMRI dataset and selected the first N complete subjects, who had completed the 7 cognitive tasks with 2 runs. The tested sample size ranges from 14 to 1060 subjects. The time window was fixed as 10s for this test.

### Saliency map of graph convolutions

In addition to high classification accuracy, good interpretability is also very important for brain decoding. In our case, we need to map which discriminative features in the brain help to differentiate different cognitive task conditions. There are several ways to visualize a deep neural network, including visualizing layer activation (Springenberg et al., 2014) and filters (Olah et al., 2017), and heatmaps of class activation (Selvaraju et al., 2017). Here, we chose the first method due to its easy implementation and generalization to graph convolutions. The basic idea is that if an input is relevant, a little variation on it will cause high change in the layer activation. This can be characterized by the gradient of the output given the input, with the positive gradients indicating that a small change to the input signals increases the output value. To visualize the gradients, we could simply use a backward pass of the activation of a single unit through the network. However, this type of map is usually very noisy, and uninversible pooling operations and nonlinear activation functions can bias the gradient. To alleviate these problems, Springenberg and his colleagues proposed to suppress the flow of gradients through neurons wherein either of input or incoming gradients were negative (Springenberg et al., 2014). Specifically, for the graph signal *X*^*l*^ of layer *l* and its gradient *R*^*l*^, the overwritten gradient 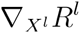 can be calculated as follows:

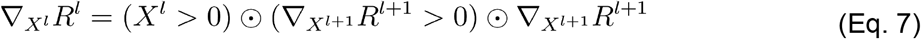

In order to generate the saliency map, we started from the output layer of a pre-trained model and used the above chain rule to propagate the gradients at each layer until reaching the input layer. This guided-backpropagation approach can provide a high-resolution saliency map which has the same dimension as the input data. Since we have used multiple time channels in the first layer of the GCN model, the approach also provides one saliency map per time step. We further calculated a heatmap of saliency maps by taking the variance across the time steps for each parcel. Since each task condition can evoke different shapes of hemodynamic response, the variance of the saliency curve provides a simplified way to evaluate the contribution of task-evoked hemodynamic response. This saliency value was additionally normalized to the range [0,1], with its highest value at 1 (a dominant effect for task prediction) and lowest at 0 (no contribution to task prediction). Note that the saliency maps were generated by using the decoding model trained from a single cognitive domain with a time window as long as the event trials.

The stability of saliency maps was evaluated among different trials, between different subjects, and at different temporal resolutions. Three different decoding models were pertrained using BOLD signals at different time windows, e.g. 0-12s, 6-12s and 0-6s within each motor task trial. For each time window, we generated the saliency maps of the motor task using 24 subjects from the test set, including 2 functional runs for each subject, 10 motor trials within each run, in total of 480 individual task trials and consequently 480 saliency maps (one saliency map per trial). The variability of salience values was analyzed not only at the subject level, by calculating the mean and std among 20 individual trials for each subject, but also at the group level, by plotting the values among 24 subjects. Three regions were selected to evaluate the inter-trial and inter-subject variability, including region a (labelled as “area 5m” in the Glasser atlas) selectively activated during foot movements, region b (labelled as “area 1”) selectively activated during hand movements, regions c (labelled as “area OP4”) selectively activated during tongue movements.

## Results

### State annotation using Brain Graph Convolutional Networks (GCN)

#### Cognitive states can be decoded with high accuracy from 10s of fMRI activity

The GCN state annotation model (Fig 1) was evaluated using the cognitive battery of HCP task-fMRI dataset acquired from 1200 healthy subjects. ***The entire dataset was split into training (70%), validation (10%) and test (20%) sets at the subject level, such that the model was evaluated on the validation set at the end of each training epoch and the best model over 100 epochs, which achieved the highest accuracy on the validation set, was saved and further evaluated on the independent test set***. During model training and evaluation, fMRI response to different cognitive tasks acquired in HCP was collected and input to the decoding model at the same time. In our study we focused on 21 task conditions spanning six cognitive domains, namely: emotion, language, motor, relational, social, and working memory. The detailed description of these cognitive tasks can be found in (Barch et al., 2013) and is also summarized in Table 1. Using a 10-second window of fMRI time series, the 21 conditions can be identified with an average test accuracy of ***90% (mean=90*.*7%, std=0*.*2% by using 10 fold cross-validation with shuffle splits)***, significantly different from the chance level of 4.8%. The confusion matrix (see Fig 2), which indicates the proportion of true and false predictions given a cognitive task state, showed a nice block diagonal architecture which means the majority of the cognitive tasks were accurately identified. After summarizing the confusion matrix according to the six cognitive domains (see Fig 2-Supplement 1), each cognitive domain could be identified with an accuracy greater than 91%. Among the six cognitive domains, the language tasks (story vs math) and motor tasks (left/right hand, left/right foot and tongue) were the most recognizable conditions, and they showed the highest precision and recall scores (average f1-score = 95% and 94%, respectively for language and motor conditions).

**Fig 2.**
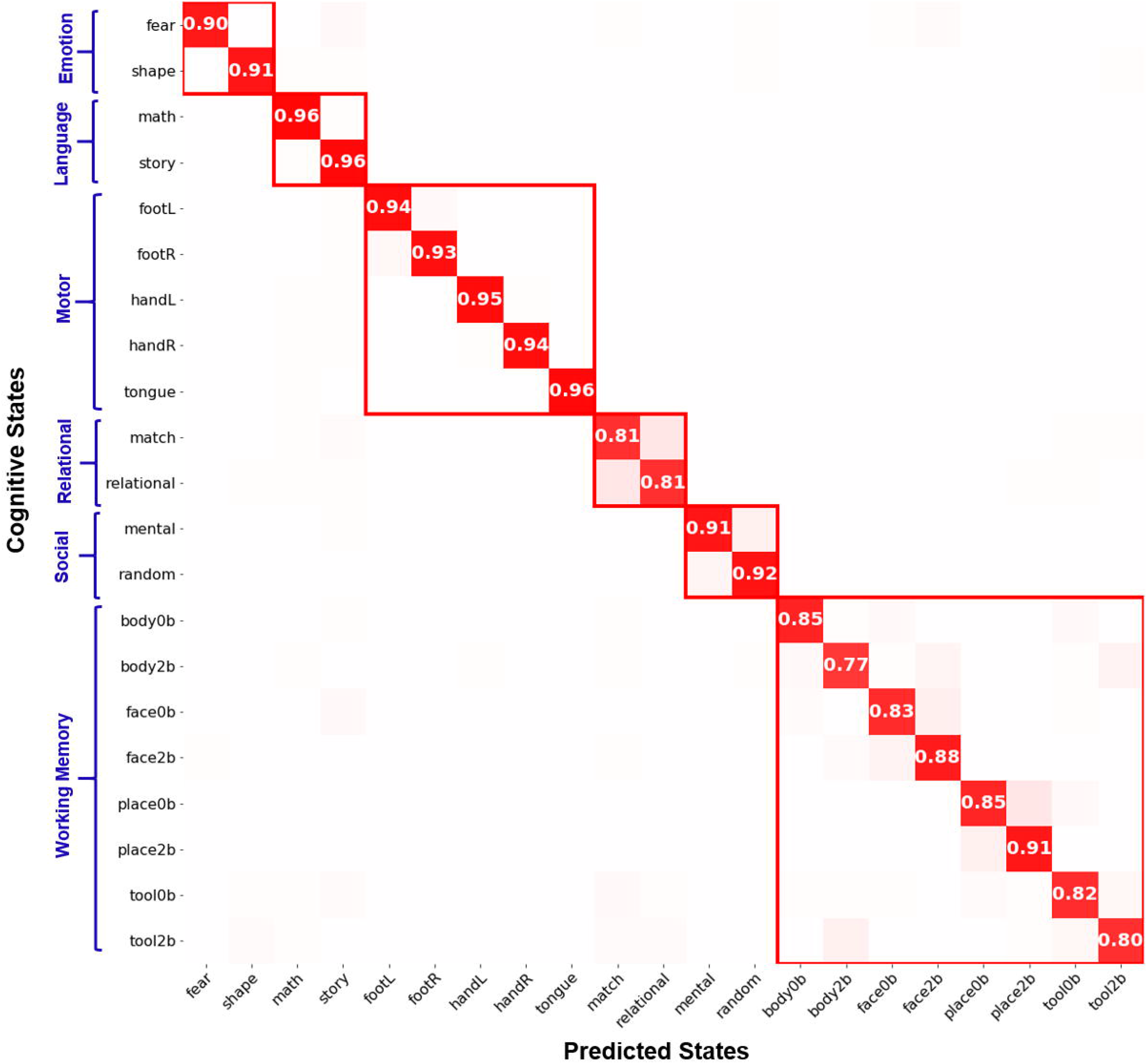
Confusion matrix of decoding 21 cognitive states. The confusion matrix was normalized by each cognitive state (row) such that each element in the matrix shows the recall score that among all predictions (column) how many of them are positive predictions. The confusion matrix showed a nice block diagonal architecture which means the majority of the cognitive tasks were accurately identified. Among the six cognitive domains, the language and motor tasks achieved the highest sensitivity, with the relational processing and working memory tasks as the lowest. Gambling task was excluded from this analysis due to the short events of the experimental design.

#### Classification errors are due to high similarity in task stimuli

Misclassifications of cognitive states not only existed within a cognitive domain but also across multiple cognitive domains. First of all, task trials within the same cognitive domain were relatively easy to be misclassified. For instance, most misclassifications of relational processing task trials were found between relational processing and pattern matching conditions. Similar misclassifications were noted between the 0-back and 2-back conditions for the working memory task (see Fig 2-Supplement 2A and B). Similar levels of false classification rates were observed when the decoding model was trained by exclusively using fMRI data from a single cognitive domain (misclassification rates as high as 13% for relational processing vs pattern-matching conditions, 10% for 0-back vs 2-back conditions). By contrast, for face and place working memory stimuli, brain decoding reached high accuracy, regardless of using a multidomain or single-domain classifier (misclassification rates less than 0.2%). This high accuracy is possibly driven by the known, strong spatial segregation of the neural representation for face vs place image, in the fusiform face area and parahippocampal place area respectively (Golarai et al., 2007). ***The reported confusion matrix somewhat resembled the representational similarity analysis in category-specific representations evoked by different visual stimuli*** *(Khaligh-Razavi and Kriegeskorte, 2014)*. Secondly, task trials can also be misclassified across different cognitive domains, probably due to similar cognitive demands of the underlying cognitive processes. For instance, we found some of the emotion and relational processing conditions were misclassified as working memory tasks. One of the reasons could be that the experimental design of the emotion task involves the matching of faces, overlapping with face encoding and retrieval in working memory tasks. Similarly, the relational processing task requires matching of drawn objects based on specific physical characteristics of target images, for instance, shape or texture, somewhat resembling the encoding and retrieval of bodies and tools in working memory tasks. These results suggest that the brain decoding model is mainly driven by the cognitive demands of the tasks and may not follow the original design of hierarchical organization among cognitive domains.

#### Decoding accuracy associated with in-scanner performance

We found a strong association between the prediction accuracy of GCN state annotation and participant’s in-scanner performance, measured using the median reaction time (RT) and average accuracy (ACC) of repeated task trials (Fig 3). For instance, during relational processing task which consists of two conditions, i.e. relational processing and pattern matching, participants reacted faster to the matching condition than relational processing (mean RT=1.48s vs 2.02s, T-val=14.88, pval=3.9e-40) with higher accuracy (mean ACC=86% vs 65%, T-val=13.18, pval=3.4e-33). Similarly, GCN also achieved higher prediction for pattern matching than relational processing (mean F1-score=0.96 vs 0.91, T-val=4.24, pval=2.7e-5). Moreover, within each condition, GCN achieved higher accuracy on trials when participants were more engaged which was indicated as shorter reaction time (Spearman rank correlation rho= -0.21, pval= 0.002) and higher accuracy (rho=0.18, pval=0.012). ***No association with subjects’ age, handedness and head motion was found at either the subject level (Fig 3-Supplement 1) or at the trial level (Fig 3-Supplement 2)***.

**Fig 3.**
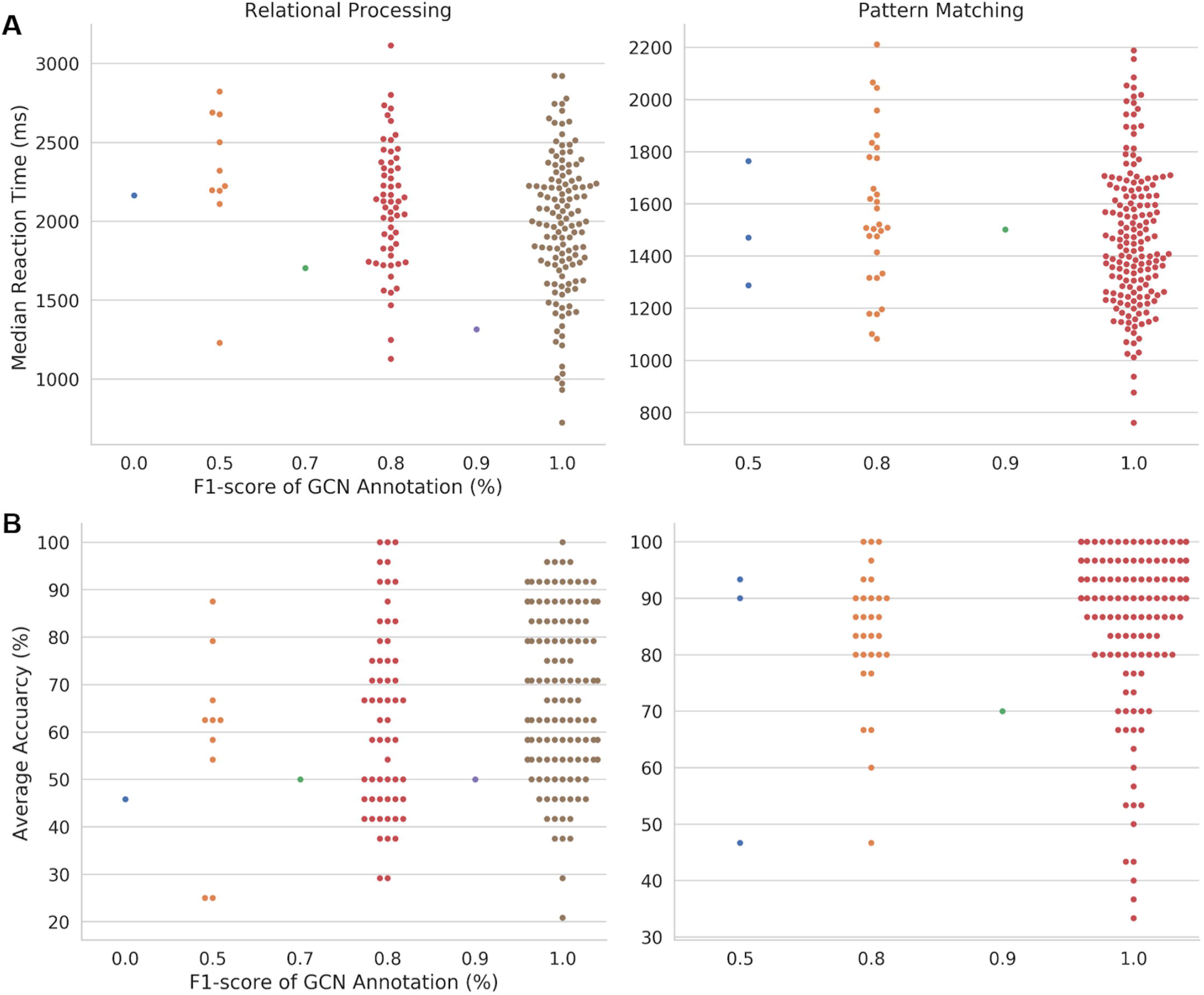
Association between prediction accuracy of GCN state annotation and participants’ in-scanner performance for all trials of the relational processing task. The relational processing task consists of two conditions, i.e. relational processing and pattern matching with each task block lasting for 16s. Two types of performance were measured for each task block, including median reaction time (A) and average accuracy (B) across repeated mini-trials. Comparing the two task conditions, participants reacted faster with higher accuracy for the pattern matching task than relational processing. Similarly, GCN also achieved higher prediction for matching (F1-score = 0.96) than relational processing (F1-score = 0.91). Within each task condition, GCN achieved higher accuracy on trials when participants responded faster (A) or achieved higher accuracy (B). The analysis was performed on 200 subjects from the test set.

#### Visualization of learned neural representations

To visualize the learned representations of cognitive functions, we projected the high-dimensional graph representations, i.e. the output of the last graph convolutional layer, onto a 2-dimensional space using t-SNE (Maaten and Hinton, 2008). We observed a high clustering effect in the learned representations (see Fig 4C). Specifically, the samples of different movement types were highly separated from each other, with the largest distance existing between tongue and foot movements. Meanwhile, the samples of the same type of movements were located closest to each other. Moderate distance was found between left and right for both hand and foot movements. A similar pattern was also observed by calculating the correlations of the learned representations across all trials (see Fig 4A and Fig 4B). But, this effect was not observed by directly projecting the input fMRI time-series or during the early stages of the training process, for which the samples from all categories collapsed into a ball (Fig 4-Supplement 1 and 2). ***Graph representations learned at each GCN layer were shown in Fig 4-Supplement 3, which indicated that the five categories of body movements were highly separable until the 5th GCN layer***.

**Fig 4.**
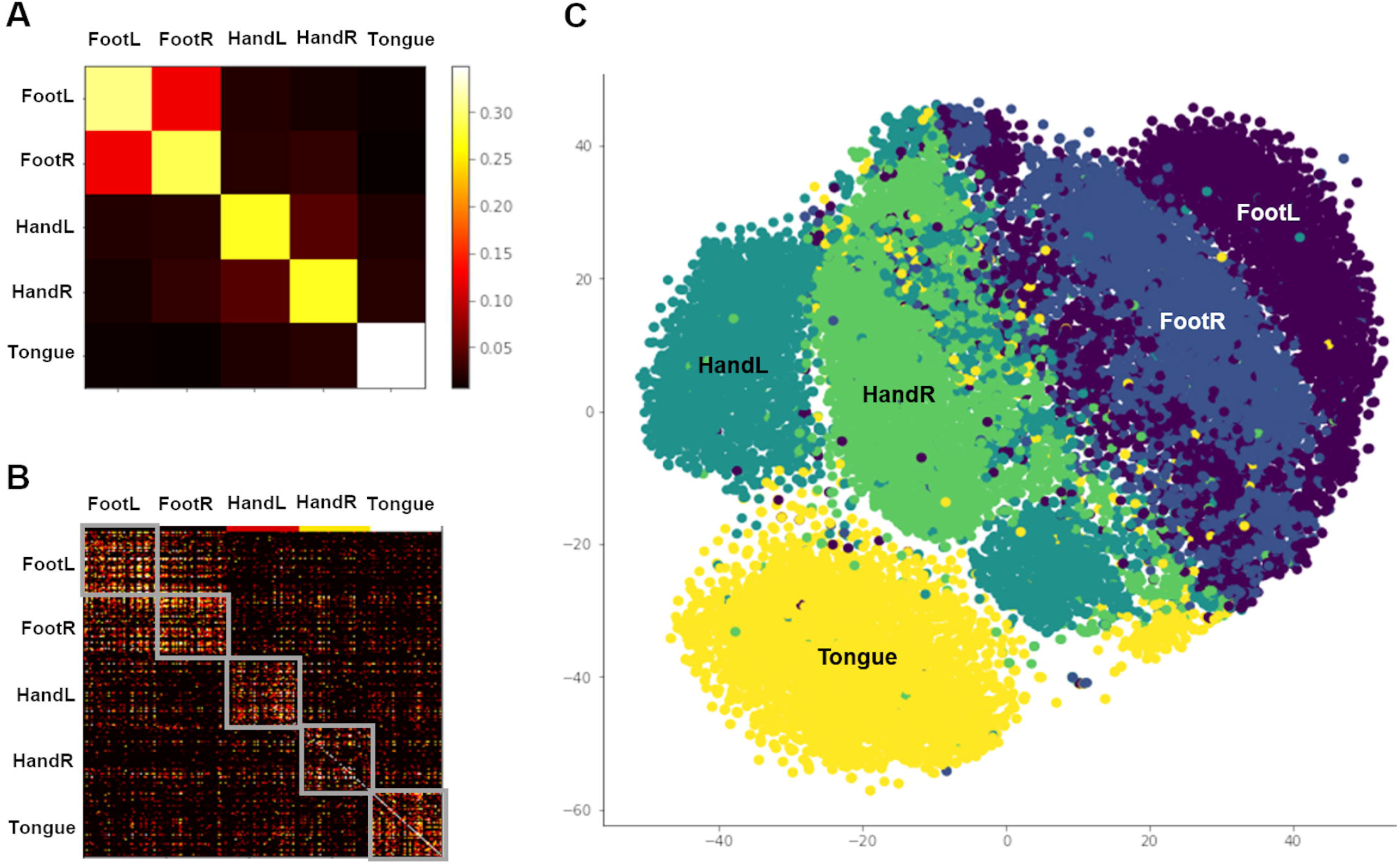
Similarity analysis of learned representations from the Motor task-fMRI data. A pre-trained single-domain GCN annotation model was used for this analysis, which meant the training set only included fMRI signals from the corresponding cognitive domain. Then, the fMRI time series from the test set was passed through the model as input and the layer activations of the last graph convolutional layer were extracted as the graph representations of brain dynamics. The similarity of graph representations was evaluated by calculating Pearson correlation between each pair of brain states (A) and experimental trials (B). Moreover, the learned representations were projected to 2-dimensional space by using t-SNE (C).

#### GCN outperformed linear and nonlinear decoding models

To establish whether the usage of deep GCN brings a substantial improvement over more traditional machine learning tools, we evaluated the same brain decoding tasks using a multi-class support vector machine classification (SVC) with ***linear and nonlinear kernels***, as our baseline model. The results showed that ***using averaged BOLD signals from every 10s of task blocks as inputs***, SVC-linear achieved much lower prediction accuracy in classifying the 21 states (90% vs 64% respectively for GCN and SVC-linear) and took longer time for training (560s vs 4221s). ***Using a nonlinear kernel, SVC-RBF achieved higher decoding accuracy (73*.*8%) but still lower than GCN. A summary of comparison between these decoding models was shown Table S1 and Fig 2-Supplement 4***. When only focusing on a single cognitive domain (using a 10s time window), SVC with linear and nonlinear kernels still showed much lower performance than GCN (87% vs 90% vs 97% for the motor task; 70% vs 77% vs 87% for working memory conditions). We also evaluated a simple multilayer perceptron (*MLP*) consisting of two hidden layers to decode brain activity over 21 states. MLP showed some improvements over the linear model, but not as high as GCN (90% vs 74% respectively for GCN and MLP). ***As an additional baseline approach, we combined multiple single-domain decoders to classify all 21 task conditions using a transfer learning strategy. Specifically, we first trained a single GCN decoder for each of the six cognitive domains. Then, the representations learned from the six single-domain decoders (i*.*e. layer activations from at the last GCN layer) were concatenated into a large feature vector. Finally, a simple classifier of MLP was trained on the concatenated features to classify all task conditions. Our results revealed that combining multiple single-domain decoders achieved lower decoding performance than directly training a multidomain decoder (83*.*6% vs 90% respectively for combining single-domain decoders and using a multidomain decoder). Besides, compared to transferring features from multiple single-domain decoders, the multi-domain decoding model not only improved the classification accuracies within each cognitive domain but mostly reduced the misclassifications between different cognitive domains (Fig 2 Supplement-3). This finding confirms that multidomain brain decoding is necessary to reliably associate spatial patterns with cognitive states and that decoders trained on a narrow set of conditions do not generalize well for other domains***.

#### Saliency maps demonstrate biologically meaningful features learned by GCN

We investigated whether GCN learns a set of biologically meaningful features during model training. For this purpose, we generated the saliency maps on the trained model by propagating the non-negative gradients backwards to the input layer (Springenberg et al., 2014). An input feature is *salient or important* only if its little variation causes big changes in the decoding output. Thus, high values in the saliency map indicate large contributions during the prediction of cognitive states. To note that the model used in this analysis was trained by exclusively using fMRI data from a single cognitive domain.

The two language conditions, story and mathematics, shared a number of salient features, likely related to shared cognitive processes. First, both conditions involve the processing of auditory statements, which may explain high salience in the primary auditory cortex and perisylvian language-related brain regions, consisting of inferior frontal gyrus (IFG), supramarginal gyrus/angular gyrus, and superior temporal gyrus (STG) (see Fig 5A). Second, the block design of both story and math conditions included a presentation and a response phase, and thus potentially imposed a high memory load on participants, and may explain the salience in the inferior parietal sulcus. There were also some salient features found only for either mathematics or story. For instance, the story condition involved salient features in more anterior part of left IFG, including pars triangularis and orbitalis. By contrast, mathematical statements involved more posterior parts, including pars opercularis of IFG and precentral sulcus. Additional inferior temporal regions were salient for mathematics only, which have been shown to be more involved in mathematical than non-mathematical judgment tasks (Amalric and Dehaene, 2016). Finally, left-lateralized salient features in IFG and STG were only revealed for the story condition, coinciding with the study showing strong lateralization for listening comprehension (Berl et al., 2010).

**Fig 5.**
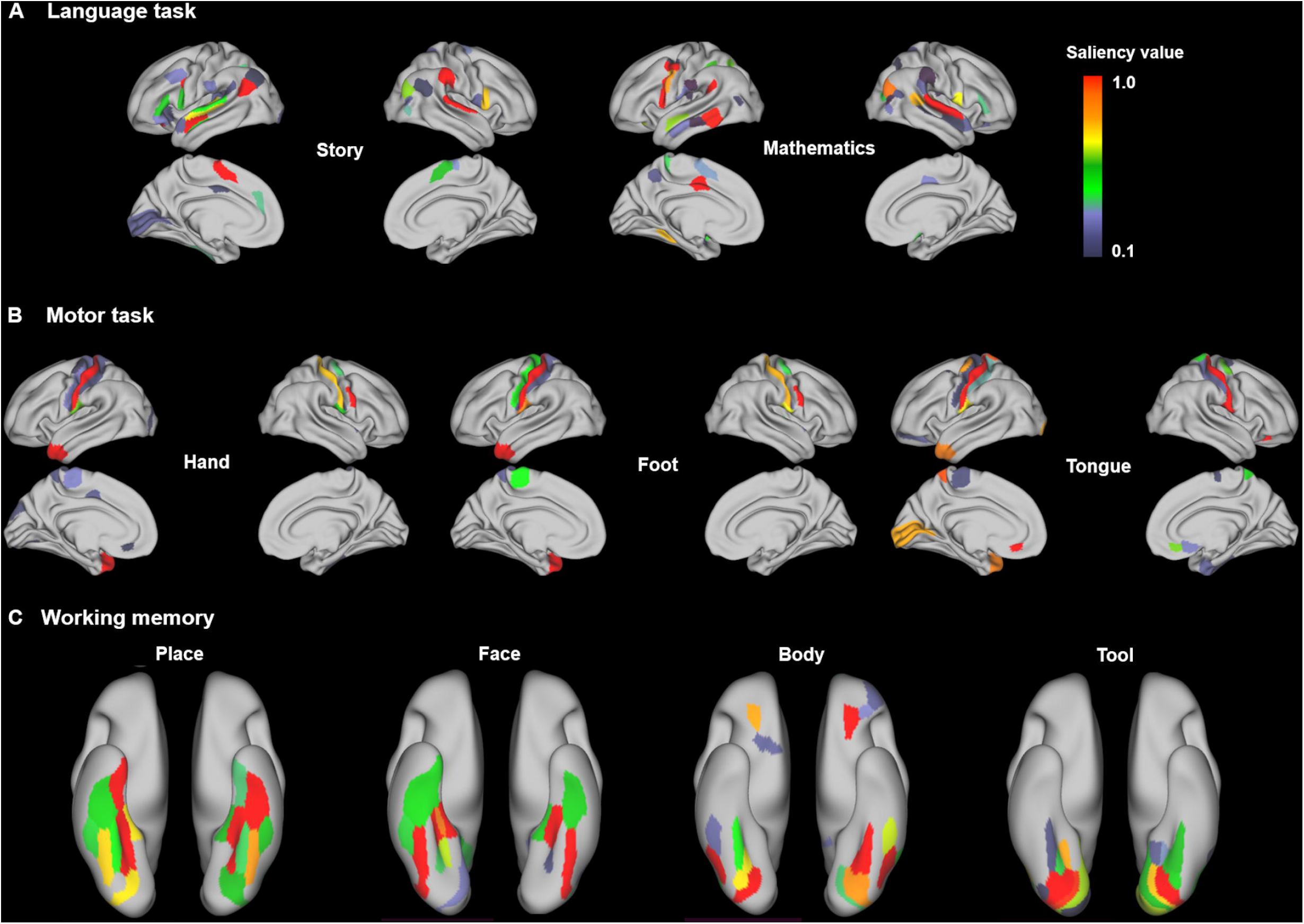
Saliency maps of language, motor and working memory tasks. (A) The story and math conditions showed high salience in the primary auditory cortex and perisylvian language-related brain regions. (B) Different types of movements were associated with salient features in the motor and somatosensory cortex. (C) The 0-back working memory task mostly engaged the ventral visual stream for encoding different types of images. The saliency maps were estimated by using the guided backpropagation based on the pre-trained single-domain GCN annotation models that only used fMRI signals from the corresponding cognitive domain during model training. A high saliency value indicates that little variation of the input features causes big changes in the decoding output. The saliency value was normalized to the range [0,1], with its highest value at 1 (a dominant effect for task prediction) and lowest at 0 (no contribution to task prediction). Only values above 0.1 were shown here to indicate a strong impact on the prediction.

As expected, no salient features in the perisylvian language-related brain regions were found for the motor task. Different types of movements were associated with high salience in the primary motor and somatosensory cortices (see Fig 5B), which have long been shown to be the main territories engaged during movements of the human body (Penfield and Boldrey, 1937). No clear somatotopic organization among different types of movements were identified here, which was somewhat expected because the primary motor and somatosensory cortex were parcellated into single strips in the Glasser’s atlas (Glasser et al., 2016). Some category-specific salient features were still identified, for instance in medial primary motor cortex for foot movement and in lateral orbitofrontal cortex for tongue movement. Unexpectedly, salient features in the left temporal pole were found for all movements. This area has been shown to support language comprehension and production (Ardila et al., 2014), which may be related to the word cues used to initiate different types of movements.

Moreover, salient features in the ventral visual stream were identified for image recognition in the working memory task (see Fig 5C). Specifically, the place stimuli activated more medial areas in the ventral temporal cortex including parahippocampal gyrus; while the face stimuli activated more lateral ventral temporal regions including fusiform gyrus. This observation is consistent with the well-known segregation of the neural substrates for encoding faces vs places, in the fusiform face area and parahippocampal place area respectively (Golarai et al., 2007).

We also found some overlap in brain regions between our saliency maps and meta-analysis activation maps from neuroquery (Fig 5-Supplement 1), as well as contrast maps from HCP dataset (Fig 5-Supplement 2). For instance, the inferior frontal gyrus and superior temporal gyrus were identified for the story condition in all three maps, while the inferior frontal sulcus and the adjacent middle frontal gyrus were identified for the mathematics condition, probably counting for the requirement of working memory for a sequence of mathematical operations. Although some consistent patterns of activations were observed for the motor tasks (Fig 5-Supplement 3), we found a large degree of divergence after mapping them onto the Glasser’s atlas (Glasser et al., 2016), probably due to the primary motor and somatosensory cortex being parcellated into single strips in the atlas and not differentiating the somatotopic areas in the feature space. For image recognition, the ventral visual stream was identified in all three maps, but with different specific spatial locations. Overall, although some overlap existed between saliency maps and meta-analysis maps, there was no systematic correspondence. This likely reflects the fact that features identified through traditional statistical tests and predictive models are to some extent divergent (Bzdok and Ioannidis, 2019). Similar observations were made with contrast maps from the HCP dataset.

***The stability analysis of the salience maps was evaluated over 480 motor trials from 24 individual subjects. We found substantial inter-trial variability (between individual trials of the same subject) and inter-subject variability in the salience values of selected regions in the motor and somatosensory cortex. Robust category-specific patterns in the salience values were also revealed, that were highly consistent across subjects (Fig 5-Supplement 4 A). For instance, region a (labelled as “area 5m” in the Glasser atlas) selectively activated during foot movements, region b (labelled as “area 1”) selectively activated during hand movements, region c (labelled as “area OP4”) selectively activated during tongue movements***.

In summary, the regions highlighted by the saliency maps are consistent with prior knowledge from the neuroscience literature, and suggest that the GCN model has learned biologically meaningful features, rather than relying on confounding effects, for example motion artifacts.

### Impact of the duration of fMRI time windows on cognitive annotations

#### Cognitive tasks showed different sensitivity levels to the duration of time windows

The temporal sensitivity of GCN was first evaluated by progressively increasing the duration of the fMRI time windows (Fig 6). At a temporal resolution of one fMRI volume (720ms), GCN could predict the 21 task conditions with an average accuracy of 56%, markedly lower than using 10 sec time windows, yet still significantly higher than chance level (4.8%). As the duration of fMRI time windows became larger, the prediction accuracy gradually increased and ***reached a peak of 90% at 10s of fMRI time series (the shortest length of task blocks among all task conditions)***. Using 6s of fMRI data, GCN already showed good performance with an average prediction accuracy of 82%. The cognitive tasks showed different levels of sensitivity to the duration of time windows ***and also plateaued at different temporal duration (Fig 6-Supplement 2)***. Among the cognitive domains, the decoding accuracy of relational processing and working memory conditions were highly dependent on the duration of time windows and required more than 10s to reach stable performance (Fig 6-Supplement 1). These domains also showed the lowest prediction accuracy for all durations of time windows. By contrast, predictions on language and social tasks reached high accuracy for durations as small as a single fMRI volume (70% and 66% for conditions of language and social tasks, respectively). This divergence on the temporal sensitivity might be driven by the form of stimuli that successive trials were used for the relational processing and working memory tasks while an auditory/video stream with continuous stimulation was presented for the language and social tasks.

**Fig 6.**
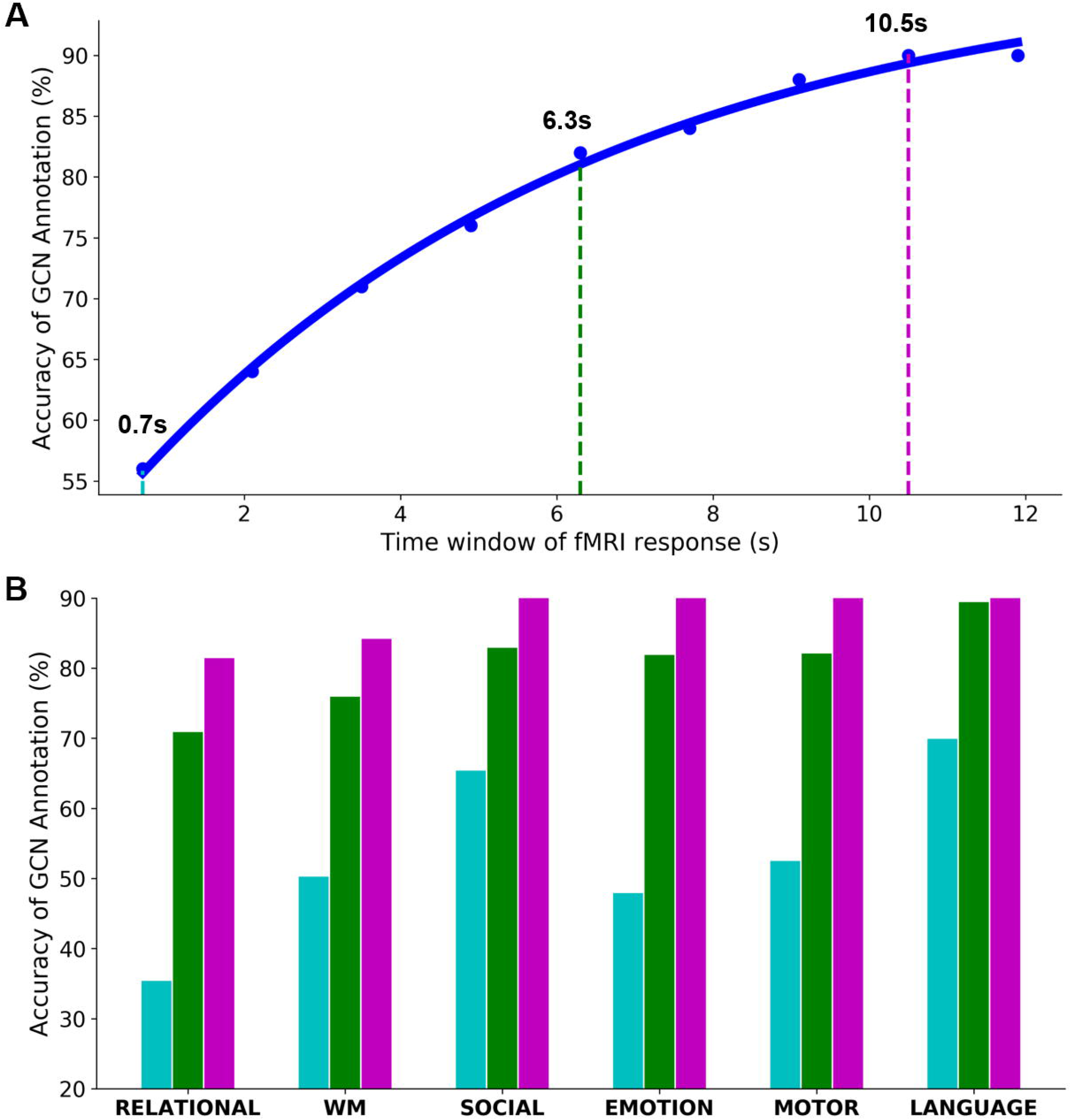
Temporal sensitivity of GCN on the fMRI time window. The temporal sensitivity of GCN was investigated with variable lengths of time windows, ranging from a single fMRI volume (0.7s) to 10s with a step of 2 TRs (i.e. 1.4s). (A) The performance of GCN annotation gradually increased as prolonging the time window of fMRI time series. It started with 56% of test accuracy on a single fMRI volume (cyan), quickly increased to 82% with 6s of fMRI data (green), and reached a plateau of 90% at 10s (purple). (B) The cognitive tasks showed high diversity in the sensitivity to the duration of time windows. Among the cognitive domains, the relational processing and working memory tasks were most sensitive to the time window and achieved the lowest decoding accuracy at all temporal scales.

#### The performance of GCN annotation is constrained by the hemodynamic response

The low performance at shorter fMRI time windows could be caused by two factors: 1) fewer parameters in decoding model especially in the first GCN layer (i.e. time window * graph filters); 2) a delay effect of the task-evoked hemodynamic response (HRF) of BOLD signals, that typically includes a dominant peak at 4-6s, and washes out around 8-12s after the end of the stimulus. To evaluate the impact of the hemodynamic response in GCN performance, we reformulated the prediction accuracy of GCN annotation on a single fMRI volume as a function of time-elapsed-from-onset. As shown in Fig 7, the GCN state annotation had an initial low performance at the cue phase, which gradually increased to reach a plateau at 6-8s after task onset. This effect was observed for all states of the motor and working memory tasks. For instance, the predictions on hand, foot and tongue movements reached an asymptotic performance of 95% for a single fMRI volume acquired 6s after task onset (Fig 7A). For the working memory task, the performance was more variable depending on the task conditions. Specifically, for the 0-back working memory task (Fig 7B), performance reached a plateau at around 8s and fluctuated around this asymptotic level. By contrast, for the 2-back working memory task (Fig 7C), the plateau was only reached at 10s after onset, and some conditions even showed a decreased performance after 20s, for example, the 2-back recognition of body and tool images.

**Fig 7.**
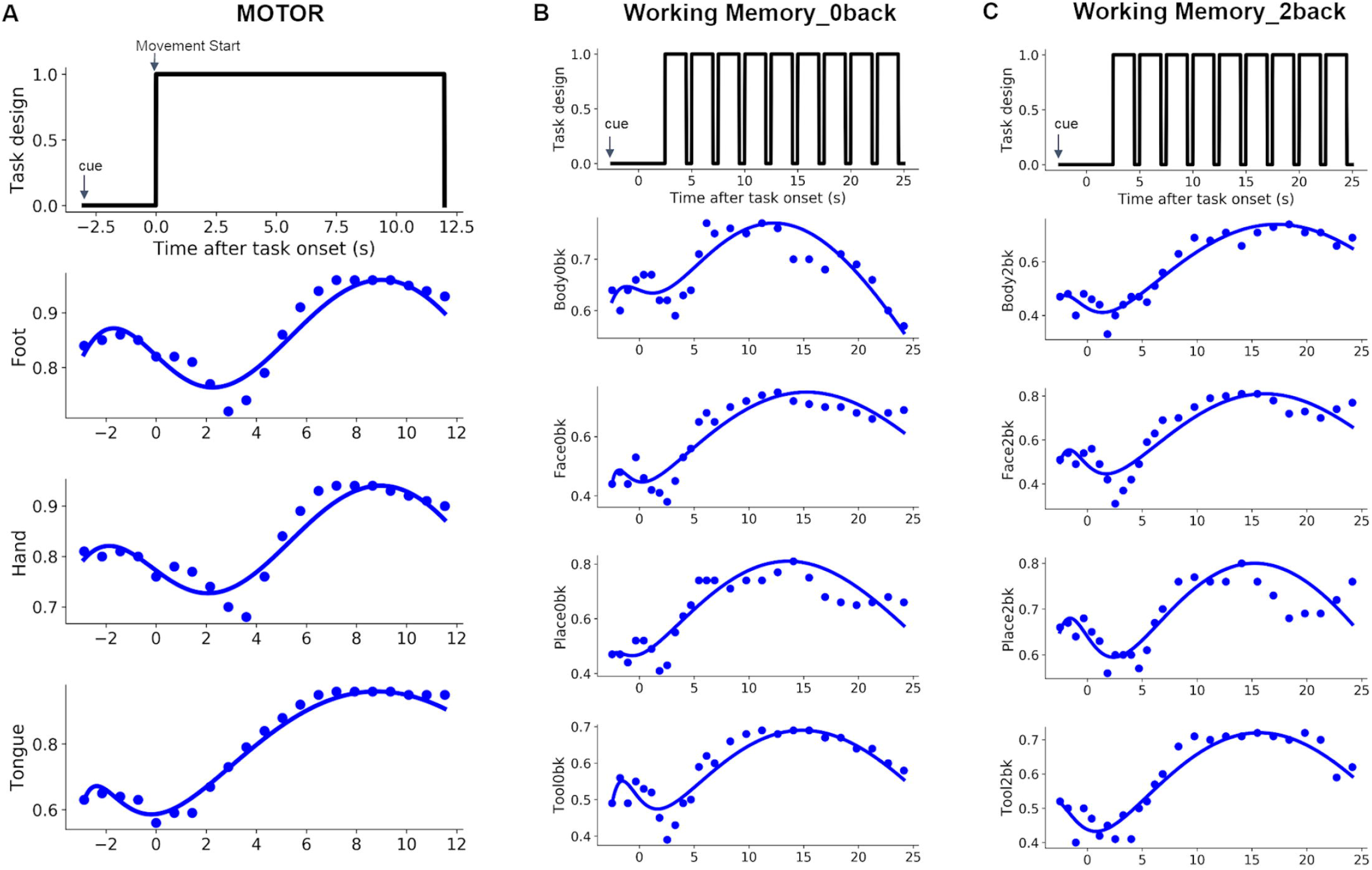
Performance of the GCN annotation as a function of time following onset. The pre-trained single-domain GCN annotation models were used for this analysis by exclusively using fMRI signals from a single cognitive domain during model training. The time window was set to 0.7s such that each single fMRI volume was treated as an independent sample. The trained model was then used to predict the cognitive state of all fMRI volumes from the test set as a function of time following onset. The state annotation of the motor (A) and working memory (B and C) tasks indicated an initial low performance at the cue phase, gradually converging to the plateau at 6-8s after the onset of a task, and then a variable post-stimulus undershoot. This resembles the effect of the task-evoked hemodynamic response of fMRI signals. Notably, GCN annotation on the motor task even achieved over 90% of test accuracy by decoding on a single fMRI volume acquired 6s after the task onset. The performance was more variable for the working memory task, e.g. lower accuracy for the 2-back conditions compared to the 0-back, but with a reverse observation for the face recognition conditions (i.e. peak performance of 75% vs 81% respectively for the 0-back and 2-back face recognition conditions).

These results suggest that for event-related designs (i.e. with short time duration of each trial), fMRI signals recorded at least 4s after the onset of the task will be required to achieve a stable GCN performance. This observation may also explain the low performance of GCN on the gambling task, where each trial only lasted 3.5s (1.5s for button press, 1s for feedback and 1s for ITI). To verify whether this rule applies to longer event trials, each task trial was split into multiple bins of 6s-time window before and after the peak of HRF. The results in Table 2 and Fig 7-Supplement 1 indicated that, with the same length of time window, GCN achieved higher performance when the BOLD signals already reached the peak of HRF, but before reaching the post-stimulus undershoot. ***The saliency maps at different time windows, e*.*g. 6-12s and 0-6s within a motor task trial, also indicated that more robust and category-specific salient features were identified by using BOLD signals after reaching the peak of HRF (Fig 7-Supplement 2)***.

**Table 2.** Performance of GCN annotation using mini-blocks of a 6s-time window before and after the peak of HRF. Task trials were split into mini-blocks with a temporal duration of 6s. Event blocks from the motor task last for 12s and thus were split into 2 mini-blocks of 6s time window. Event blocks from the working memory task last for 25s and thus were split into 4 mini-blocks of 6s time windows. These mini-blocks were treated as independent samples during model training. We also trained and evaluated separate decoding models for each of the time windows, by exclusively using the fMRI time series from the corresponding time bins. The last column indicates the average decoding performance on a mixture of 6s mini-blocks by including fMRI signals at all different phases.

| Task domain | Decoding Performance on Time windows |  |  |  |  |
| --- | --- | --- | --- | --- | --- |
|  | 0-6s | **6-12s** | 12-18s | 18-24s | Mixed (6s time window) |
| Motor | 85.51 % | **94.58 %** | N/A | N/A | 88.60 % |
| Working memory | 75.54 % | **85.72 %** | 82.59 % | 81.89 % | 80.37 % |
| ALL tasks | 79.43 % | **88.38 %** | N/A | N/A | 81.51 % |

### Impact of population sample size on cognitive annotations

#### GCN annotation reached a performance plateau with around 280 subjects

The sensitivity of GCN on sample size was investigated by changing the number of independent subjects selected from HCP task-fMRI dataset, ranging from 14 to 1060 subjects who have collected 2 sessions of all cognitive tasks. These subjects were again split into training (70%), validation (10%) and test (20%) sets. Generally, with more subjects, GCN achieved higher accuracy in decoding the 21 cognitive states (Fig 8). GCN annotation already achieved decent performance with a handful of subjects (average f1-score=46% using 14 subjects). Performance quickly increased to 77% by using 71 subjects and reached a plateau of 85% with around 280 subjects. After that, performance only showed slight improvement by using larger data samples. Different cognitive tasks showed different highly variable sensitivity to sample size, and also varied in the asymptotic performance of the model. Specifically, the relational processing and working memory required the largest sample size: 284 subjects and 213 subjects, respectively, to reach 85% of the asymptotic performance. By contrast, the language and motor tasks only required 35 and 57 individuals, respectively, to reach 85% of the asymptotic performance. This variation on the sensitivity of sample size might be caused by different levels of inter-subject variability in the cognitive demands of the underlying cognitive processes. For instance, large individual variability has been reported in working memory tasks (Fougnie et al., 2012; Osaka et al., 2003), while the language network was consistently activated during the auditory language comprehension across different populations and languages (Friederici, 2011; Wu et al., 2019; Zhang et al., 2017).

**Fig 8.**
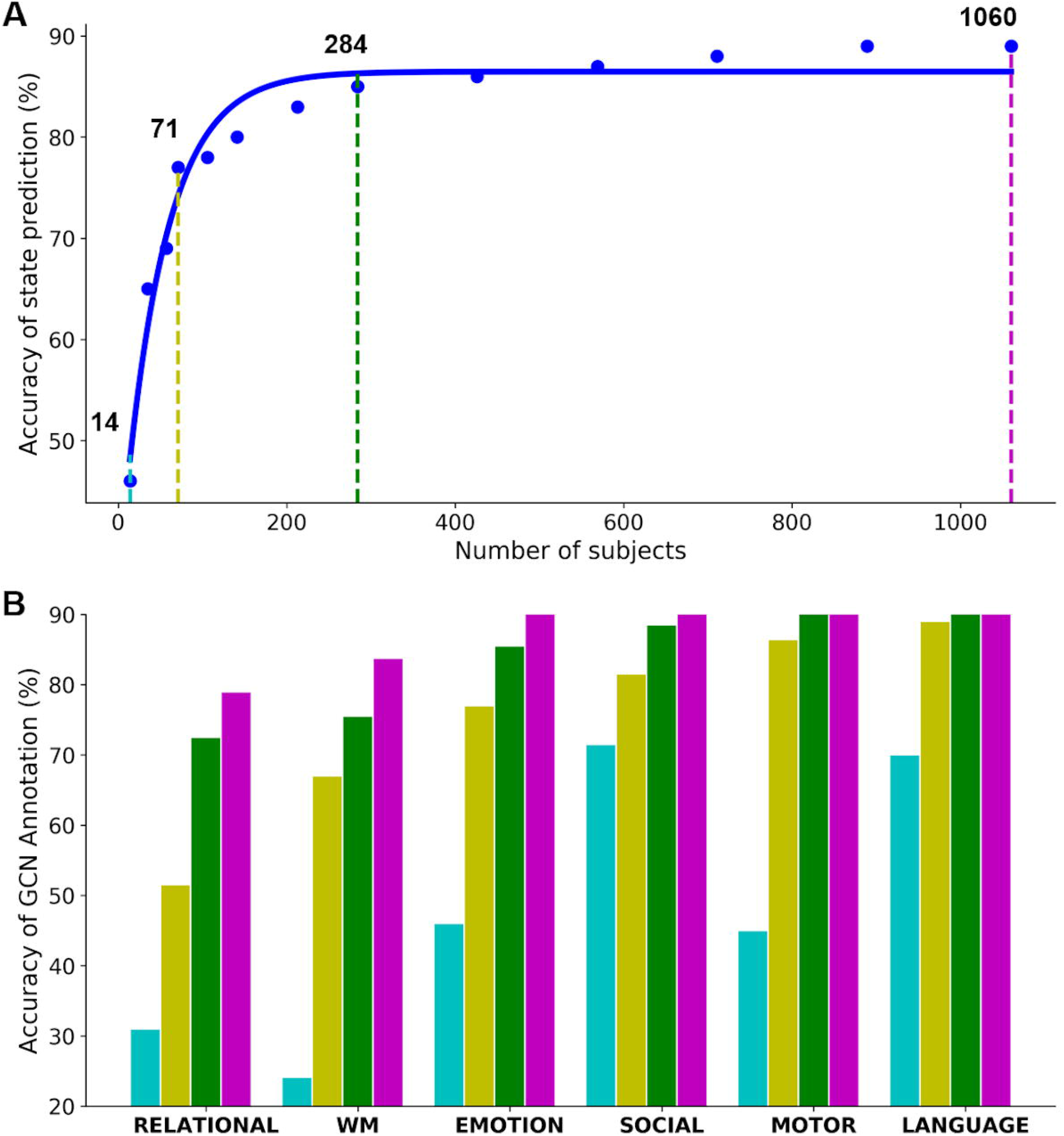
Sensitivity of GCN on sample size of independent subjects. The sensitivity of GCN on sample size (A) was investigated by changing the number of independent subjects selected from HCP task-fMRI dataset, ranging from 14 to 1060 with a smaller step before the plateau and a larger step after. GCN annotation starts with 46% of test accuracy in decoding the 21 cognitive states by using only 14 subjects (cyan). Then, the performance quickly increased to 77% by using 71 subjects (yellow) and reached a plateau of 85% with around 280 subjects (green). Among the cognitive domains (B), the relational processing and working memory were the most demanding tasks on the sample size, while the language and motor tasks were more robust to the size of the dataset.

## Discussion

The present study proposed a generalized brain decoding model which annotates brain dynamics of 21 cognitive functions using a short series of fMRI volumes. This approach relies on convolutional operations on a brain graph, which leverages our prior knowledge on network organization of human brain cognition. Graph convolution integrates information of brain dynamics among distributed brain networks and generates robust neural representations that could be generalizable across a large group of population and multiple cognitive domains. Specifically, our model identified 21 experimental conditions across 6 cognitive domains simultaneously with an accuracy of 90% on unseen subjects, by only using 10s window of fMRI signals. This high performance on brain annotation was mainly contributed by brain response of biologically meaningful brain areas, in line with the literature on functional localizers for each cognitive domain, as revealed by the saliency maps. By examining variable time windows, we found that our decoding model achieved above-chance annotation with a fine temporal resolution, as short as a single fMRI volume. Volume-to-volume performance followed the shape of a hemodynamic response, with a high accuracy achieved after at least 6 seconds following stimulus onset. Besides, the model converged to its stable performance by using a subset of 280 subjects. Together, our results provide an automated tool to annotate brain activity with fine temporal resolution and fine cognitive granularity, as well as high generalizability to new subjects.

### Decoding brain cognitive process from a short time series

Brain decoding has been a popular topic in neuroscience literature for decades. The majority of studies still focused on the recognition of visual stimuli (Haxby, 2001; Haxby et al., 2014; Huth et al., 2012). To build a decoding model that could generalize beyond visual stimuli and incorporate multiple cognitive domains is still a challenging topic. Researchers have attempted to tackle the issue of multidomain brain decoding by using meta-analytic approaches based on thousands of reported brain coordinates (Bartley et al., 2018) and a set of statistical contrast maps (Rubin et al., n.d.) from a series of published studies. But meta-analyses bring other types of limitations, such as unbalanced samples across different cognitive domains (Alamolhoda et al., 2017), publication bias towards positive effects (Dubben and Beck-Bornholdt, 2005), as well as over-estimated effect sizes from small studies (Lin, 2018). These factors may bias the decoding analysis by falsely inferring the mental states given limited available studies (see discussion in (Lieberman et al., 2016; Lieberman and Eisenberger, 2015; Wager et al., 2016)).

To avoid these biases, an alternative approach has also been proposed by training linear classifiers on the activation maps collected from a group of individuals that have been scanned over a variety of cognitive tasks (Bzdok et al., 2016; Poldrack et al., 2009; Varoquaux et al., 2018). It is worth noting that, by using parametric modelling and averaging brain response across multiple trials and even multiple runs, it is possible to achieve high accuracy on the task of distinguishing different experimental conditions, for instance classifying a subset of HCP tasks (Bzdok et al., 2016). The challenge here is to achieve such high accuracy using a fully data-driven approach to infer cognitive states directly from a short time series. This requires the decoding model to take into account not only the overall discriminative patterns of brain response under different cognitive tasks, but also their temporal dynamics, i.e. changes of brain activations over time. Such brain dynamics are usually revealed by electro- or magnetoencephalography (Kietzmann et al., 2019), but has also recently been investigated in fMRI studies. For instance, Gonzalez-Castillo and his colleagues reported distinct shapes of task-evoked hemodynamic responses among distributed brain networks during a discrimination task of letters and numbers (Gonzalez-Castillo et al., 2012). Similar findings were also revealed in our previous study that premotor and sensorimotor cortex showed different time courses during the preparation and execution stage of a motor sequential task (Orban et al., 2015). These studies suggest that the differences in the shapes of hemodynamic response can help to distinguish different conditions or stages of cognitive process. Accumulated evidence suggests that it is feasible to infer cognitive states directly from a short time window of hemodynamic response. Early attempts in the field include the reconstruction of visual scenes and prediction of semantic context from natural movies by fitting a linear regression model for fMRI signals for each individual voxel (Huth et al., 2012; Nishimoto et al., 2011). These studies neglected the modular, and hierarchical nature of brain organization by treating each brain voxel independently.

***Most existing decoding models were based on a linear classifier*** *(Haxby, 2001; Haxby et al*., ***2014; Haynes et al***., ***2007) or a linear regression model*** *(Huth et al., 2012; Mitchell et al., 2008;* ***Varoquaux et al***., ***2018). These linear models have low capacity in handling the complex and neurovascular coupled neural activity measured by BOLD signals. Our results also indicated a significant gain in decoding performance (9% boost) by replacing the linear kernel with a nonlinear kernel (e*.*g. RBF) in SVC (Fig 2-Supplement 3). In order to decode brain activity directly from a short fMRI time series, we need to use more complex and nonlinear decoding models. These decoding models are required to incorporate the high-dimensional spatiotemporal dynamics of brain responses that are shared among distributed brain networks***. Recently, promising results on brain decoding have been shown by using deep artificial neural networks (DNNs). For instance, multiple cognitive domains can be distinguished by applying convolutional neural networks on the whole-brain hemodynamic response (Wang et al., 2019). But the temporal dependence of hemodynamic response was interrupted by choosing random time points from the entire fMRI scan. This effect can be corrected by applying a recurrent neural network to brain activity instead. Li and Fan proposed a long short-term memory (LSTM) architecture to predict the cognitive states from fMRI time-series of a set of functional networks (Li and Fan, 2019). ***This recurrent network architecture achieved remarkable gains in the decoding power by taking into account the temporal dependency within task-evoked brain activity. At the same time, the model usually required relatively long time windows from the history in order to predict the next time point, for instance using 40TRs in*** *(Li and Fan, 2019)*. ***It is challenging to decode over a very short time-series down to a single fMRI volume***. In this study, we extend this line of work by combining the graph Laplacian with the DNN architecture and proposed a generalized brain decoding model that takes into account both the network architecture of the human brain (in space) and the fluctuations in the task-evoked BOLD signals (in time).

### Brain decoding using graph convolution network

Graph Laplacian provides a powerful tool to map the intrinsic organization of the human brain, including parcellating brain areas (Fan et al., 2016; Johansen-Berg et al., 2004), identifying functional areas and networks (Atasoy et al., 2016; Craddock et al., 2012), and generating connectivity gradients (Margulies et al., 2016). This approach works not only on static brain connectome but also on dynamic brain signals that fluctuate over time. Recently, studies have shown that graph Laplacian captured different modes of brain dynamics by decomposing the task-evoked BOLD signals into different frequencies (Ortega et al., 2018). Convergent evidence suggests that the low frequency modes, which have similar brain signals within a local community, corresponded to the low-level functions that localized within certain brain regions, such as motor learning (Huang et al., 2016). On the other hand, the high frequency modes, which indicate high variational signals across brain networks, were associated with high-level cognitions that distributed among multiple brain systems, such as cognitive switch (Medaglia et al., 2018). We generalized this approach by automatically learning a linear combination of the graph modes across multiple frequencies through graph convolutions, i.e. convolving the input fMRI signals with a graph filter. The resultant decoding model not only represented low-level functions like movements of body parts, but also embedded the high-level cognitions such as N-back working memory and language comprehension. The results showed that the decoding model achieved high classification accuracies on these cognitive tasks (Fig 2 and Fig 2-Supplement 1). ***Besides, the confusion matrix indicated that the misclassifications not only existed among different task conditions within the same domain but also across cognitive domains (Fig 2-Supplement 2), for instance between working memory and relational processing tasks. A similar pattern of inter-domain confusion has been reported in an independent study by exploring the spatial similarity of brain activation maps across cognitive tasks (Pinho et al***., ***2018). This misclassification was partially due to the similarity in task stimuli, for instance, the relational processing task requires matching of drawn objects based on specific physical characteristics of target images, for instance, shape or texture, somewhat resembling the encoding and retrieval of bodies and tools in working memory tasks***. Moreover, the saliency maps indicated that the task inference was drawn from brain response of biological meaningful brain regions, for instance, the motor and somatosensory cortex for the motor task, and the perisylvian language network for the story and math auditory statements (Fig 5), consistent with known brain anatomy and function (Friederici, 2011; Penfield and Boldrey, 1937).

Graph convolutions generated a new representation of brain activity by integrating neural dynamics from interconnected brain regions. A variety of neural representations were generated by training multiple graph filters at each GCN layer. Specifically, ***at the first GCN layer, various shapes of hemodynamic responses were captured and different representations of brain activity were learned by fitting different kernel weights for each time point after task onset (Fig 1-Supplement 2)***. After stacking several GCN layers, high-level graph representations were generated that integrated neural dynamics not only within specific brain networks but also across multiple networks, and even distributed across the whole brain. Our results demonstrated that the generated graph representations already include task-specific information that discriminates different experimental conditions, for instance, showing the largest distances among different types of movements, moderate distance between left and right movements, and a small distance between the same type of movements (Fig 4). Moreover, a strong association was found between the model performance on classification of graph representations and human performance on recognition of visual patterns, e.g. reaction time of relational processing and pattern matching trials in scanner (Fig 3). A similar finding has been reported previously that the high-frequency graph mode was strongly associated with the response time of trials in a cognitive switch task (Medaglia et al., 2018).

Token together, using brain graph convolutional networks, our decoding model generates high-level neural representations from brain dynamics and provides a possible solution towards multidomain brain decoding by learning various shapes of hemodynamic response and integrating such neural dynamics among multiple brain systems. ***Such representations can be used to predict the transition of brain states during cognitive processes or even at rest. For example, we applied the pretrained multidomain decoding model to predict the cognitive states using resting-state fMRI data and found that body movements and mind-wandering (i*.*e. watching clips of randomly moving shapes) were among the most frequent brain states during resting-state (Fig 7-Supplement 2). The significance of these findings would need to be further validated, using direct probes of cognitive states experienced by participants*** *(Smallwood and Andrews-Hanna, 2013)*.

### Temporal resolution of brain decoding

The temporal resolution of brain decoding has been mostly ignored in previous studies, by either using meta-analytic approaches (Bartley et al., 2018; Rubin et al., n.d.), or training classifiers on activation maps (Poldrack et al., 2009; Varoquaux et al., 2018). The recent work of Loula and colleagues (2018) demonstrated the feasibility of decoding stimuli with short inter-stimuli intervals. Temporal resolution is thus becoming an important factor for brain annotation, especially when we tried to infer cognitive functions directly from brain response. A series of impressive work has been done in Gallant’s group, in which the authors used brain response to reconstruct the visual frames of natural movies (Nishimoto et al., 2011) or to map more abstract concepts of visual objects, e.g. semantic context (Huth et al., 2012). But these studies did not directly attempt to characterize what amount of temporal data is required to perform meaningful brain decoding. The temporal resolution of fMRI decoding is intrinsically constrained by two factors, including the acquisition time for a whole-brain fMRI scan (i.e. TR) and the delay effect of hemodynamic response. With a common setting as 2 second, the acquisition time was pushed down to a third (TR = 720ms) in HCP database by using simultaneous multislice acquisitions (Uğurbil et al., 2013), which brings opportunities to investigate fine-grained temporal dynamics of brain activity.

Using this dataset, Li and Fan successfully predicted the entire experimental design of the working memory task by using a sliding window approach (Li and Fan, 2019). But each time window still took around 30s of fMRI signals as input for task inference. To which extent of a shorter time window the decoding model can work with is still unexplored. In this study, we applied graph convolutions on a short series of fMRI signals and investigated the temporal resolution of brain decoding by using variable time windows of fMRI scans, ranging from a single fMRI volume to the entire event trial. Leveraging the fast fMRI acquisition of HCP database, our model can annotate 21 cognitive conditions with a sub-second temporal resolution. In the meantime, the decoding performance was still impacted by the task-evoked hemodynamic response, for instance, higher decoding accuracy by using fMRI signals after the peak of HRF than before the peak. This phenomenon was observed not only for low-level functions like body movements, but also high-level cognitions such as working memory tasks, or even missing all experimental conditions together (Table 2 and Fig 6). ***In addition, a similar effect in the saliency maps was also observed that more stable saliency values were generated when using the time window before the peak of HRF, including lower inter-trial and inter-subject variability in the saliency values along with higher distinctions between different task conditions (Fig 5-Supplement 4)***. There are still a lot of challenges before achieving real-time brain decoding, for instance, to decode fast events with a short duration or even overlapping hemodynamic response.

### Limitations and future applications

In the current project, we only explore the block design of task-fMRI dataset, i.e. consisting of long events with repeated trials that in total last for more than 10s. However, it is still unclear how to generalize the decoding pipeline to naturalistic stimuli, for instance visual scenes from movies, which consists of short and fast-switching events. The measured BOLD signals might be a mixture of hemodynamic responses evoked by different task events. Early attempts have been made by adding independent regressors with delayed onsets (Nishimoto et al., 2011). But the simple linear model only generates a blurred image from the average prediction of each category. One possible solution to this problem is to use a multi-label decoding model based on GCN. Specifically, given a short-series of fMRI signals, the model predicts a set of cognitive states instead of one single task condition. Due to the delay effect of hemodynamic response that reaches plateau around 6s past stimulus, we can modify the label matrix by prolonging each event duration until 8s after the task onset and allow multiple labels assigned to the same time point.

***In this project, we used resting-state functional connectivity to construct a functional brain graph that was used as the fixed graph architecture in GCN. A series of studies in the literature have illustrated a great potential of using resting-state functional connectivity in predicting cognitive functions, not only in behavior*** *(Rosenberg et al., 2020)* ***but also in task-evoked brain activity*** *(Cole et al., 2016; Tavor et al., 2016)*. ***Some researchers have also applied anatomical connectivity to separate specific components in brain dynamics that were associated with high-level cognitions*** *(Huang et al., 2018;* ***Medaglia et al***., ***2018). In our next project, we will systematically investigate the impact of using different brain graphs on GCN models, for instance using a spatial graph, built based on spatial adjacency on the cortical surface, a functional graph, calculated using resting-state functional connectivity, as well as an anatomical graph derived from diffusion tractography***.

An interesting potential application of our work would be transfer learning. In natural image processing, it is common practice to take a model already trained on a large dataset, such as AlexNet trained on ImageNet (Krizhevsky et al., 2012), and fine-tune the parameters of the trained model to accomplish a new task (Tajbakhsh et al., 2016). This allows training complex models even in the absence of extensive training data. This problem of lacking a sufficiently large dataset for specific experimental questions is pervasive in medical imaging. Our model was made publicly available (https://github.com/zhangyu2ustc/GCN_fmri_decoding) and can be used as a reference model for domain adaptation, possibly making contributions in a variety of domains, including neurological and psychiatric disorders. It could also be applied in samples where extensive data is acquired on a few subjects, such as the individual brain charting (IBC) project (Pinho et al., 2018) or the Courtois project on neuronal modelling (neuromod, https://docs.cneuromod.ca).

## Acknowledgment

This work was supported in part by the Courtois foundation through the Courtois NeuroMod Project and the IVADO Postdoctoral Scholarships Program. PB is supported by a salary award of “Fonds de recherche du Québec - Santé”, chercheur boursier junior 2.

## References

Alamolhoda, M., Ayatollahi, S.M.T., Bagheri, Z., 2017. A comparative study of the impacts of unbalanced sample sizes on the four synthesized methods of meta-analytic structural equation modeling. BMC Res. Notes 10, 446.

Amalric, M., Dehaene, S., 2016. Origins of the brain networks for advanced mathematics in expert mathematicians. Proc. Natl. Acad. Sci. U. S. A.

Ardila, A., Bernal, B., Rosselli, M., 2014. The Elusive Role of the Left Temporal Pole (BA38) in Language: A Preliminary Meta-Analytic Connectivity Study. International Journal of Brain Science 2014. https://doi.org/10.1155/2014/946039

Atasoy, S., Donnelly, I., Pearson, J., 2016. Human brain networks function in connectome-specific harmonic waves. Nat. Commun. 7, 10340.

Barch, D.M., Burgess, G.C., Harms, M.P., Petersen, S.E., Schlaggar, B.L., Corbetta, M., Glasser, M.F., Curtiss, S., Dixit, S., Feldt, C., Nolan, D., Bryant, E., Hartley, T., Footer, O., Bjork, J.M., Poldrack, R., Smith, S., Johansen-Berg, H., Snyder, A.Z., Van Essen, D.C., WU-Minn HCP Consortium, 2013. Function in the human connectome: task-fMRI and individual differences in behavior. Neuroimage 80, 169–189.

Bartley, J.E., Boeving, E.R., Riedel, M.C., Bottenhorn, K.L., Salo, T., Eickhoff, S.B., Brewe, E., Sutherland, M.T., Laird, A.R., 2018. Meta-analytic evidence for a core problem solving network across multiple representational domains. Neurosci. Biobehav. Rev. 92, 318–337.

Berl, M.M., Duke, E.S., Mayo, J., Rosenberger, L.R., Moore, E.N., VanMeter, J., Ratner, N.B., Vaidya, C.J., Gaillard, W.D., 2010. Functional anatomy of listening and reading comprehension during development. Brain Lang. 114, 115–125.

Bronstein, M.M., Bruna, J., LeCun, Y., Szlam, A., Vandergheynst, P., 2017. Geometric Deep Learning: Going beyond Euclidean data. IEEE Signal Process. Mag. 34, 18–42.

Bruna, J., Zaremba, W., Szlam, A., LeCun, Y., 2013. Spectral Networks and Locally Connected Networks on Graphs. arXiv [cs.LG].

Bullmore, E., Sporns, O., 2009. Complex brain networks: graph theoretical analysis of structural and functional systems. Nat. Rev. Neurosci. 10, 186–198.

Buxton, R.B., Wong, E.C., Frank, L.R., 1998. Dynamics of blood flow and oxygenation changes during brain activation: the balloon model. Magn. Reson. Med. 39, 855–864.

Bzdok, D., Ioannidis, J.P.A., 2019. Exploration, Inference, and Prediction in Neuroscience and Biomedicine. Trends Neurosci. 42, 251–262.

Bzdok, D., Varoquaux, G., Grisel, O., Eickenberg, M., Poupon, C., Thirion, B., 2016. Formal Models of the Network Co-occurrence Underlying Mental Operations. PLoS Comput. Biol. 12, e1004994.

Cole, M.W., Ito, T., Bassett, D.S., Schultz, D.H., 2016. Activity flow over resting-state networks shapes cognitive task activations. Nat. Neurosci. 19, 1718–1726.

Craddock, R.C., James, G.A., Holtzheimer, P.E., 3rd, Hu, X.P., Mayberg, H.S., 2012. A whole brain fMRI atlas generated via spatially constrained spectral clustering. Hum. Brain Mapp. 33, 1914–1928.

Defferrard, M., Bresson, X., Vandergheynst, P., 2016. Convolutional Neural Networks on Graphs with Fast Localized Spectral Filtering. arXiv [cs.LG].

Dubben, H.-H., Beck-Bornholdt, H.-P., 2005. Systematic review of publication bias in studies on publication bias. BMJ 331, 433–434.

Fan, L., Li, H., Zhuo, J., Zhang, Y., Wang, J., Chen, L., Yang, Z., Chu, C., Xie, S., Laird, A.R., Fox, P.T., Eickhoff, S.B., Yu, C., Jiang, T., 2016. The Human Brainnetome Atlas: A New Brain Atlas Based on Connectional Architecture. Cereb. Cortex 26, 3508–3526.

Fougnie, D., Suchow, J.W., Alvarez, G.A., 2012. Variability in the quality of visual working memory. Nat. Commun. 3, 1229.

Friederici, A.D., 2011. The brain basis of language processing: from structure to function. Physiol. Rev. 91, 1357–1392.

Glasser, M.F., Coalson, T.S., Robinson, E.C., Hacker, C.D., Harwell, J., Yacoub, E., Ugurbil, K., Andersson, J., Beckmann, C.F., Jenkinson, M., Smith, S.M., Van Essen, D.C., 2016. A multi-modal parcellation of human cerebral cortex. Nature 536, 171–178.

Glasser, M.F., Sotiropoulos, S.N., Wilson, J.A., Coalson, T.S., Fischl, B., Andersson, J.L., Xu, J., Jbabdi, S., Webster, M., Polimeni, J.R., Van Essen, D.C., Jenkinson, M., WU-Minn HCP Consortium, 2013. The minimal preprocessing pipelines for the Human Connectome Project. Neuroimage 80, 105–124.

Golarai, G., Ghahremani, D.G., Whitfield-Gabrieli, S., Reiss, A., Eberhardt, J.L., Gabrieli, J.D.E., Grill-Spector, K., 2007. Differential development of high-level visual cortex correlates with category-specific recognition memory. Nat. Neurosci. 10, 512–522.

Gonzalez-Castillo, J., Hoy, C.W., Handwerker, D.A., Robinson, M.E., Buchanan, L.C., Saad, Z.S., Bandettini, P.A., 2015. Tracking ongoing cognition in individuals using brief, whole-brain functional connectivity patterns. Proc. Natl. Acad. Sci. U. S. A. 112, 8762–8767.

Gonzalez-Castillo, J., Saad, Z.S., Handwerker, D.A., Inati, S.J., Brenowitz, N., Bandettini, P.A., 2012. Whole-brain, time-locked activation with simple tasks revealed using massive averaging and model-free analysis. Proc. Natl. Acad. Sci. U. S. A. 109, 5487–5492.

Haxby, J.V., 2001. Distributed and Overlapping Representations of Faces and Objects in Ventral Temporal Cortex. Science. https://doi.org/10.1126/science.1063736

Haxby, J.V., Connolly, A.C., Guntupalli, J.S., 2014. Decoding neural representational spaces using multivariate pattern analysis. Annu. Rev. Neurosci. 37, 435–456.

Haynes, J.-D., Sakai, K., Rees, G., Gilbert, S., Frith, C., Passingham, R.E., 2007. Reading Hidden Intentions in the Human Brain. Current Biology. https://doi.org/10.1016/j.cub.2006.11.072

Huang, W., Bolton, T.A.W., Medaglia, J.D., Bassett, D.S., Ribeiro, A., Van De Ville, D., 2018. A Graph Signal Processing Perspective on Functional Brain Imaging. Proc. IEEE 106, 868–885.

Huang, W., Goldsberry, L., Wymbs, N.F., Grafton, S.T., Bassett, D.S., Ribeiro, A., 2016. Graph Frequency Analysis of Brain Signals. IEEE J. Sel. Top. Signal Process. 10, 1189–1203.

Huth, A.G., Nishimoto, S., Vu, A.T., Gallant, J.L., 2012. A continuous semantic space describes the representation of thousands of object and action categories across the human brain. Neuron 76, 1210–1224.

Johansen-Berg, H., Behrens, T.E.J., Robson, M.D., Drobnjak, I., Rushworth, M.F.S., Brady, J.M., Smith, S.M., Higham, D.J., Matthews, P.M., 2004. Changes in connectivity profiles define functionally distinct regions in human medial frontal cortex. Proc. Natl. Acad. Sci. U. S. A. 101, 13335–13340.

Khaligh-Razavi, S.M., Kriegeskorte, N., 2014. Deep supervised, but not unsupervised, models may explain IT cortical representation. PLoS Comput. Biol.

Kietzmann, T.C., Spoerer, C.J., Sörensen, L., Cichy, R.M., Hauk, O., Kriegeskorte, N., 2019. Recurrence required to capture the dynamic computations of the human ventral visual stream. arXiv preprint arXiv:1903. 05946.

Kipf, T.N., Welling, M., 2016. Semi-Supervised Classification with Graph Convolutional Networks. arXiv [cs.LG].

Krizhevsky, A., Sutskever, I., Hinton, G.E., 2012. ImageNet Classification with Deep Convolutional Neural Networks, in: Pereira, F., Burges, C.J.C., Bottou, L., Weinberger, K.Q. (Eds.), Advances in Neural Information Processing Systems 25. Curran Associates, Inc., pp. 1097–1105.

Lieberman, M.D., Burns, S.M., Torre, J.B., Eisenberger, N.I., 2016. Reply to Wager et al.: Pain and the dACC: The importance of hit rate-adjusted effects and posterior probabilities with fair priors. Proc. Natl. Acad. Sci. U. S. A.

Lieberman, M.D., Eisenberger, N.I., 2015. The dorsal anterior cingulate cortex is selective for pain: Results from large-scale reverse inference. Proc. Natl. Acad. Sci. U. S. A. 112, 15250–15255.

Li, H., Fan, Y., 2019. Interpretable, highly accurate brain decoding of subtly distinct brain states from functional MRI using intrinsic functional networks and long short-term memory recurrent neural networks. NeuroImage. https://doi.org/10.1016/j.neuroimage.2019.116059

Lin, L., 2018. Bias caused by sampling error in meta-analysis with small sample sizes. PLoS One 13, e0204056.

Maas, A.L., Hannun, A.Y., Ng, A.Y., 2013. Rectifier nonlinearities improve neural network acoustic models, in: Proc. Icml. p. 3.

Maaten, L. van der Hinton, G., 2008. Visualizing Data using t-SNE. J. Mach. Learn. Res. 9, 2579–2605.

Margulies, D.S., Ghosh, S.S., Goulas, A., Falkiewicz, M., Huntenburg, J.M., Langs, G., Bezgin, G., Eickhoff, S.B., Castellanos, F.X., Petrides, M., Jefferies, E., Smallwood, J., 2016. Situating the default-mode network along a principal gradient of macroscale cortical organization. Proc. Natl. Acad. Sci. U. S. A. 113, 12574–12579.

Medaglia, J.D., Huang, W., Karuza, E.A., Kelkar, A., Thompson-Schill, S.L., Ribeiro, A., Bassett, D.S., 2018. Functional Alignment with Anatomical Networks is Associated with Cognitive Flexibility. Nat Hum Behav 2, 156–164.

Mensch, A., Mairal, J., Bzdok, D., Thirion, B., Varoquaux, G., 2017. Learning Neural Representations of Human Cognition across Many fMRI Studies. arXiv [stat.ML].

Mitchell, T.M., Shinkareva, S.V., Carlson, A., Chang, K.-M., Malave, V.L., Mason, R.A., Just, M.A., 2008. Predicting human brain activity associated with the meanings of nouns. Science 320, 1191–1195.

Nishimoto, S., Vu, A.T., Naselaris, T., Benjamini, Y., Yu, B., Gallant, J.L., 2011. Reconstructing visual experiences from brain activity evoked by natural movies. Curr. Biol. 21, 1641–1646.

Olah, C., Mordvintsev, A., Schubert, L., 2017. Feature visualization. Distill 2, e7.

Orban, P., Doyon, J., Petrides, M., Mennes, M., Hoge, R., Bellec, P., 2015. The Richness of Task-Evoked Hemodynamic Responses Defines a Pseudohierarchy of Functionally Meaningful Brain Networks. Cereb. Cortex 25, 2658–2669.

Ortega, A., Frossard, P., Kovacevic, J., Moura, J.M.F., Vandergheynst, P., 2018. Graph Signal Processing: Overview, Challenges, and Applications. Proc. IEEE 106, 808–828.

Osaka, M., Osaka, N., Kondo, H., Morishita, M., Fukuyama, H., Aso, T., Shibasaki, H., 2003. The neural basis of individual differences in working memory capacity: an fMRI study. Neuroimage 18, 789–797.

Penfield, W., Boldrey, E., 1937. Somatic motor and sensory representation in the cerebral cortex of man as studied by electrical stimulation. Brain 60, 389–443.

Pinho, A.L., Amadon, A., Ruest, T., Fabre, M., Dohmatob, E., Denghien, I., Ginisty, C., Becuwe-Desmidt, S., Roger, S., Laurier, L., Joly-Testault, V., Médiouni-Cloarec, G., Doublé, C., Martins, B., Pinel, P., Eger, E., Varoquaux, G., Pallier, C., Dehaene, S., Hertz-Pannier, L., Thirion, B., 2018. Individual Brain Charting, a high-resolution fMRI dataset for cognitive mapping. Sci Data 5, 180105.

Poldrack, R.A., 2011. Inferring mental states from neuroimaging data: from reverse inference to large-scale decoding. Neuron 72, 692–697.

Poldrack, R.A., 2006. Can cognitive processes be inferred from neuroimaging data? Trends Cogn. Sci. 10, 59–63.

Poldrack, R.A., Halchenko, Y.O., Hanson, S.J., 2009. Decoding the large-scale structure of brain function by classifying mental States across individuals. Psychol. Sci. 20, 1364–1372.

Powers, D.M., 2011. Evaluation: from precision, recall and F-measure to ROC, informedness, markedness and correlation.

Raj, A., Kuceyeski, A., Weiner, M., 2012. A network diffusion model of disease progression in dementia. Neuron 73, 1204–1215.

Raj, A., LoCastro, E., Kuceyeski, A., Tosun, D., Relkin, N., Weiner, M., 2015. Network Diffusion Model of Progression Predicts Longitudinal Patterns of Atrophy and Metabolism in Alzheimer’s Disease. Cell Reports. https://doi.org/10.1016/j.celrep.2014.12.034

Rosenberg, M.D., Scheinost, D., Greene, A.S., Avery, E.W., Kwon, Y.H., Finn, E.S., Ramani, R., Qiu, M., Constable, R.T., Chun, M.M., 2020. Functional connectivity predicts changes in attention observed across minutes, days, and months. Proc. Natl. Acad. Sci. U. S. A. 117, 3797–3807.

Rubin, T.N., Koyejo, O., Gorgolewski, K.J., Jones, M.N., Poldrack, R.A., Yarkoni, T., n.d. Decoding brain activity using a large-scale probabilistic functional-anatomical atlas of human cognition. https://doi.org/10.1101/059618

Selvaraju, R.R., Cogswell, M., Das, A., Vedantam, R., Parikh, D., Batra, D., 2017. Grad-cam: Visual explanations from deep networks via gradient-based localization, in: Proceedings of the IEEE International Conference on Computer Vision. pp. 618–626.

Shuman, D.I., Narang, S.K., Frossard, P., Ortega, A., Vandergheynst, P., 2013. The emerging field of signal processing on graphs: Extending high-dimensional data analysis to networks and other irregular domains. IEEE Signal Process. Mag. 30, 83–98.

Smallwood, J., Andrews-Hanna, J., 2013. Not all minds that wander are lost: the importance of a balanced perspective on the mind-wandering state. Front. Psychol. 4, 441.

Springenberg, J.T., Dosovitskiy, A., Brox, T., Riedmiller, M., 2014. Striving for Simplicity: The All Convolutional Net. arXiv [cs.LG].

Tajbakhsh, N., Shin, J.Y., Gurudu, S.R., Hurst, R.T., Kendall, C.B., Gotway, M.B., Jianming Liang, 2016. Convolutional Neural Networks for Medical Image Analysis: Full Training or Fine Tuning? IEEE Trans. Med. Imaging 35, 1299–1312.

Tavor, I., Jones, O.P., Mars, R.B., Smith, S.M., 2016. Task-free MRI predicts individual differences in brain activity during task performance.

Ugurbil, K., Xu, J., Auerbach, E.J., Moeller, S., Vu, A.T., Duarte-Carvajalino, J.M., Lenglet, C., Wu, X., Schmitter, S., Van de Moortele, P.F., Strupp, J., Sapiro, G., De Martino, F., Wang, D., Harel, N., Garwood, M., Chen, L., Feinberg, D.A., Smith, S.M., Miller, K.L., Sotiropoulos, S.N., Jbabdi, S., Andersson, J.L.R., Behrens, T.E.J., Glasser, M.F., Van Essen, D.C., Yacoub, E., WU-Minn HCP Consortium, 2013. Pushing spatial and temporal resolution for functional and diffusion MRI in the Human Connectome Project. Neuroimage 80, 80–104.

Varoquaux, G., Schwartz, Y., Poldrack, R.A., Gauthier, B., Bzdok, D., Poline, J.-B., Thirion, B., 2018. Atlases of cognition with large-scale human brain mapping. PLOS Computational Biology. https://doi.org/10.1371/journal.pcbi.1006565

Wager, T.D., Atlas, L.Y., Botvinick, M.M., Chang, L.J., Coghill, R.C., Davis, K.D., Iannetti, G.D., Poldrack, R.A., Shackman, A.J., Yarkoni, T., 2016. Pain in the ACC? Proceedings of the National Academy of Sciences 113, E2474–E2475.

Wang, X., Liang, X., Jiang, Z., Nguchu, B.A., Zhou, Y., Wang, Y., Wang, H., Li, Y., Zhu, Y., Wu, F., Gao, J.-H., Qiu, B., 2019. Decoding and mapping task states of the human brain via deep learning. Hum. Brain Mapp. https://doi.org/10.1002/hbm.24891

Wu, C.-Y., Zaccarella, E., Friederici, A.D., 2019. Universal neural basis of structure building evidenced by network modulations emerging from Broca’s area: The case of Chinese. Hum. Brain Mapp. 40, 1705–1717.

Zhang, Y., Fan, L., Caspers, S., Heim, S., Song, M., Liu, C., Mo, Y., Eickhoff, S.B., Amunts, K., Jiang, T., 2017. Cross-cultural consistency and diversity in intrinsic functional organization of Broca’s Region. Neuroimage 150, 177–190.

