## Supplemental figures for "Functional Annotation of Human Cognitive States using Deep Graph Convolution"

### Materials and Methods

#### fMRI Datasets

The HCP task protocols were listed below and also listed in Table 2. More detailed description of the tasks can be found in (Barch et al., 2013).

##### Motor task

Participants are presented with visual cues that ask them to either tap their fingers, or squeeze toes, or move the tongue. Each block of a movement type (hand, foot or tongue) is preceded by a 3s cue and lasts for 12s. In each of the two runs, there are 13 blocks in total, including 2 blocks of tongue movements, 4 of hand movements and 4 of foot movements, as well as 3 additional fixation blocks (15s) in the middle of each run.

##### Language task

The language task consists of two conditions, i.e. story or mathematics, with variable duration of auditory statements. During the story trials, participants listen to brief auditory stories (5-9 sentences) adapted from Aesop's fables, followed by a two-alternative-choice question and response on the topic of the story. In the math trials, participants are presented with a series of arithmetic operations, e.g. addition and subtraction, followed by a two-alternative-choice question and response about the result of the operations. The math task is adaptive to maintain a similar level of difficulty across participants. Overall, the mathematical trials lasts around 12-15 seconds while the story trials lasts 25-30 seconds.

##### Working memory task

The working memory task involves two-levels of cognitive functions, with a combination of the category recognition task and N-Back memory task. Specifically, participants are presented with pictures of places, tools, faces and body parts. These 4 different stimulus types are presented in separate blocks, with half of the blocks using a 2-back working memory task (showing the same image after two image blocks) and the other half using a 0-back working memory task (showing the same image in the next block). Each of the two runs contains 8 task blocks and 4 fixation blocks (15s). Each task block consists of a 2.5s cue indicating the task type, followed by 10 task trials (2.5s each). For each task trial, the stimulus is presented for 2 seconds, followed by a 500 ms inter-task interval (ITI) when participants need to respond as target or not.

##### Social Cognition task

Participants are presented with short video clips of objects (squares, circles, triangles) that either interacted in some way, or moved randomly on the screen. After each video clip, participants need to judge whether the objects had a mental interaction , Not Sure, or No interaction. Each of the two runs contains 5 video blocks (20s) and 5 fixation blocks (15s). There are equal length of video blocks between the types of conditions among the 2 task runs  (2 Mental and 3 Random in run 1, 3 Mental and 2 Random in run 2)

##### Relational Processing task

The task consists of two conditions, i.e. relational processing and matching. In the relational processing condition, participants are presented with 2 pairs of objects, which are shown in 6 different shapes and filled with 6 different textures. Participants need to first decide whether the top pair of objects differ in shape or texture and then make the final decision whether the bottom pair differ along that same dimension. Each relational block consists of 4 task trials, where the stimuli are presented for 3500 ms followed by a 500 ms ITI. In the control matching condition, only one top pair of objects and one bottom object are presented. Additionally, the matching dimension is specified by a cue word presented in the middle of the screen (either “shape” or “texture”). Participants need to decide whether the bottom object matches either of the top objects on that dimension. Each matching block consists of 5 task trials, where the stimuli are presented for 2800 ms followed by a 400 ms ITI. In each of the two runs, there are 3 relational blocks, 3 matching blocks and 3 fixation blocks (16s). Each task block lasts 16 seconds.

##### Emotion Processing

The task consists of two conditions, i.e. face or shape images. Participants need to match the two images presented on the bottom of the screen to the target image  whether the image shown at the top of the screen. The face images can have either an angry or fearful expression. In each of the two runs, there are 3 face blocks, 3 shape blocks and 1 fixation block (8s) at the end of each run. Each task block is preceded by a 3s task cue indicating the task type (“shape” or “face”), followed by 6 task trials (3s each). For each task trial, the stimulus is presented for 2 seconds, followed by a 1-second ITI when participants need to respond to which of the bottom images matches the target.

### Results

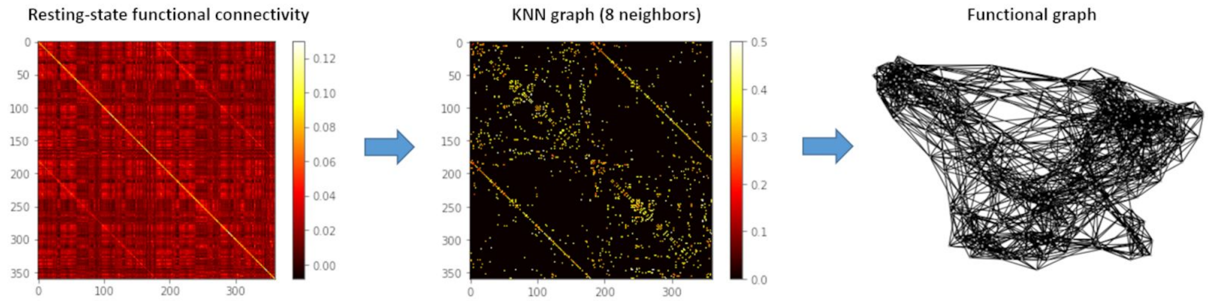


**Fig 1-Supplement 1. The illustration of constructing a functional brain graph.**

First, group averaged resting-state functional connectivity (RSFC) were calculated based on minimal preprocessed resting-state fMRI data from $N = 1080$ HCP subjects. Then, a k-nearest-neighbour (k-NN) graph was built by only connecting each brain region to its 8 neighbours with highest (positive) connectivitions. The resulting brain graph was visualized using networkx (https://networkx.github.io/).

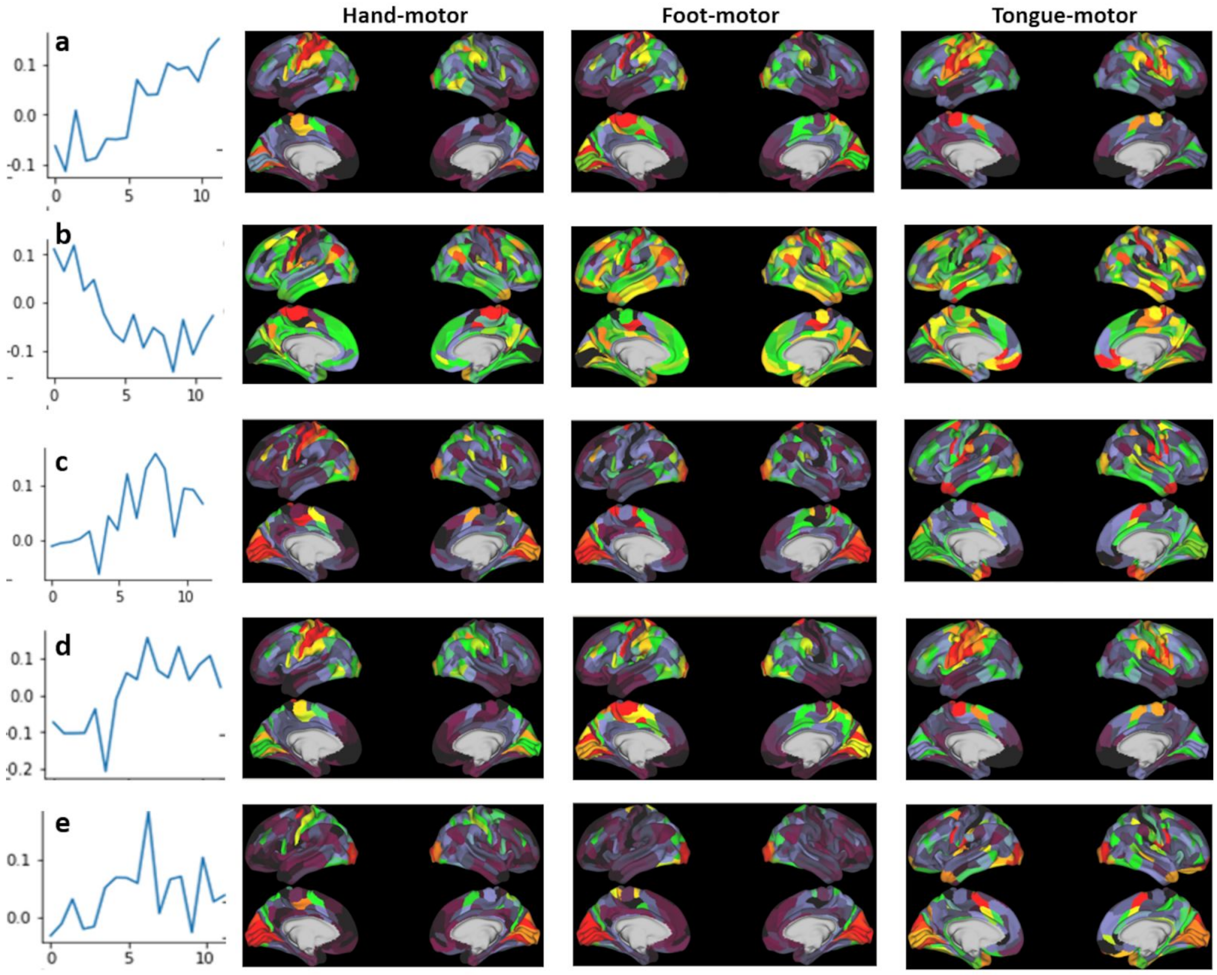


**Fig 1-Supplement 2. Visualization of temporal filters learned in the first GCN layer and their corresponding activation patterns in the brain.**

During model training, the first GCN layer learns various versions of convolution kernels for fMRI time-series, as replacement of the canonical hemodynamic response function (HRF). The shapes of temporal convolutional kernels were shown in the first column. And the corresponding "activation patterns" for each movement category were shown in the second-to-fourth columns. Generally speaking, when assigning positive weights to the time window 5-10s and negative weights to 0-5s (for instance, a and d), the model detected strong activation maps in the motor and sensory cortex along with category-specific representations for each movement type, for instance high activations in medial primary motor cortex for foot movement and in lateral orbitofrontal cortex for tongue movement. By contrast, when using a reverse convolutional kernel by assigning negative weights to 5-10s (for instance, b), the model detected a uniform activation pattern at the whole-brain level with low task specificity in representations. Other shapes of convolutional kernels (e.g. c and e) were learned to detect localized activation patterns that showing universal representations across all movements, for instance, detecting strong activation patterns in visual cortex (e).

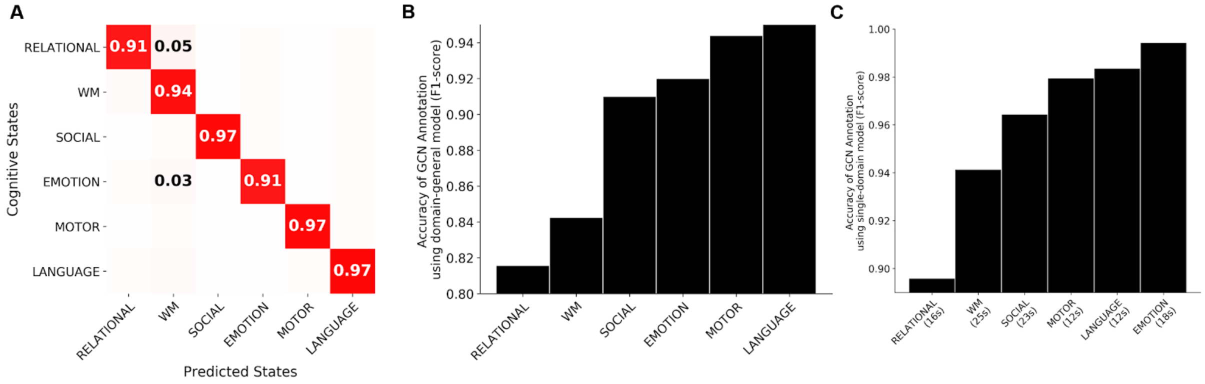


**Fig 2-Supplement 1. Confusion matrix and F1-scores of the six cognitive domains.**

The normalized confusion matrix (A) indicates the sensitivity of each cognitive domain by averaging the recall score within each of the six domains. Relational processing and working memory showed the lowest sensitivity, with some misclassifications between emotion/relational processing and working memory tasks. A similar trend was shown in the F1-scores of GCN annotation using the decoding model either trained on multiple domains simultaneously (B) or exclusively using a single domain (C). Both of them showed the highest decoding accuracy for language and motor tasks and the lowest for relational processing and working memory tasks. Comparing the two models, a significant improvement of prediction accuracy was also shown for all cognitive domains.

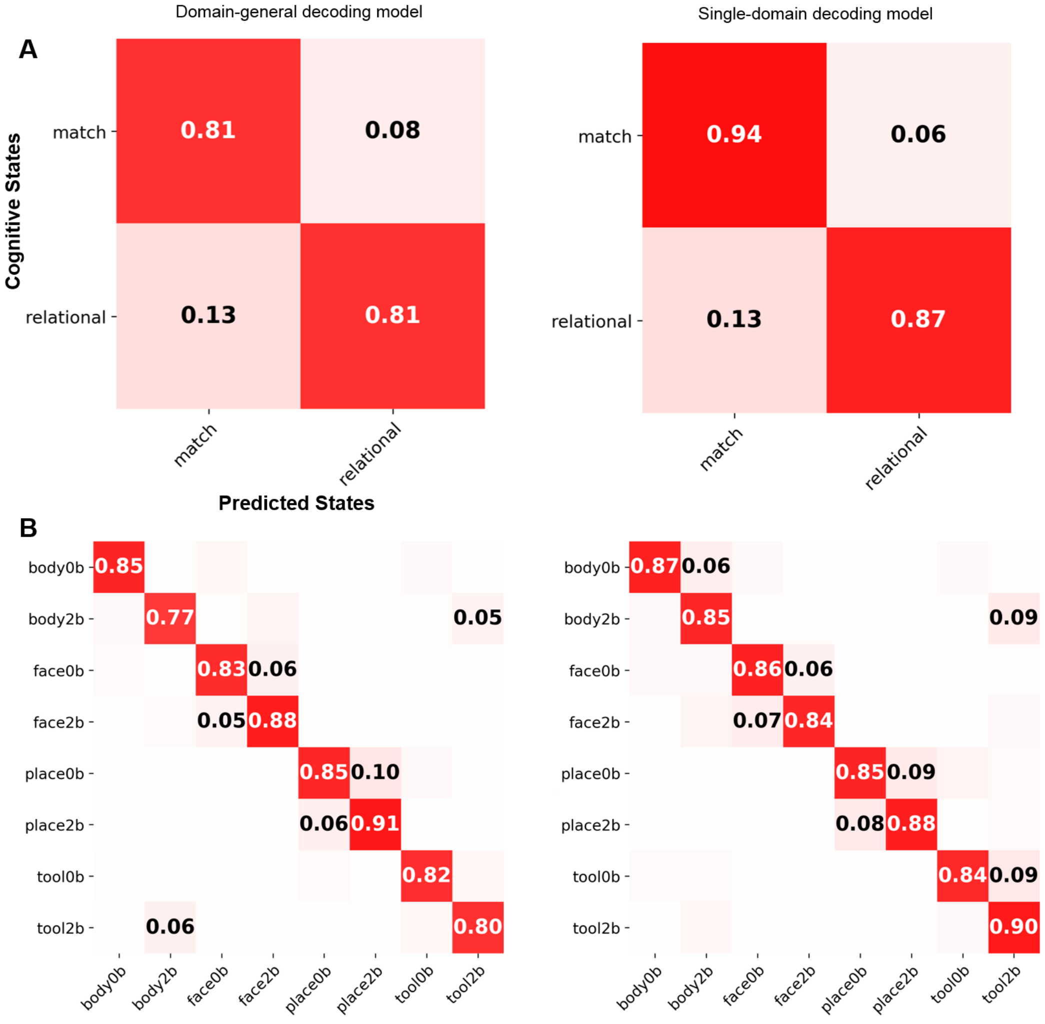


**Fig 2-Supplement 2. Confusion matrices for the relational processing (A) and working memory tasks (B).**

The confusion matrix was either extracted from the domain-general decoding model which encodes 21 cognitive conditions simultaneously (left panel) or calculated using a separate decoding model for every single cognitive domain (right panel). ALL decoding models were trained using 10s of fMRI time series. A similar level of misclassification rates was found for the two types of decoding models, with a slight improvement of prediction accuracy for the model trained exclusively from a single domain.

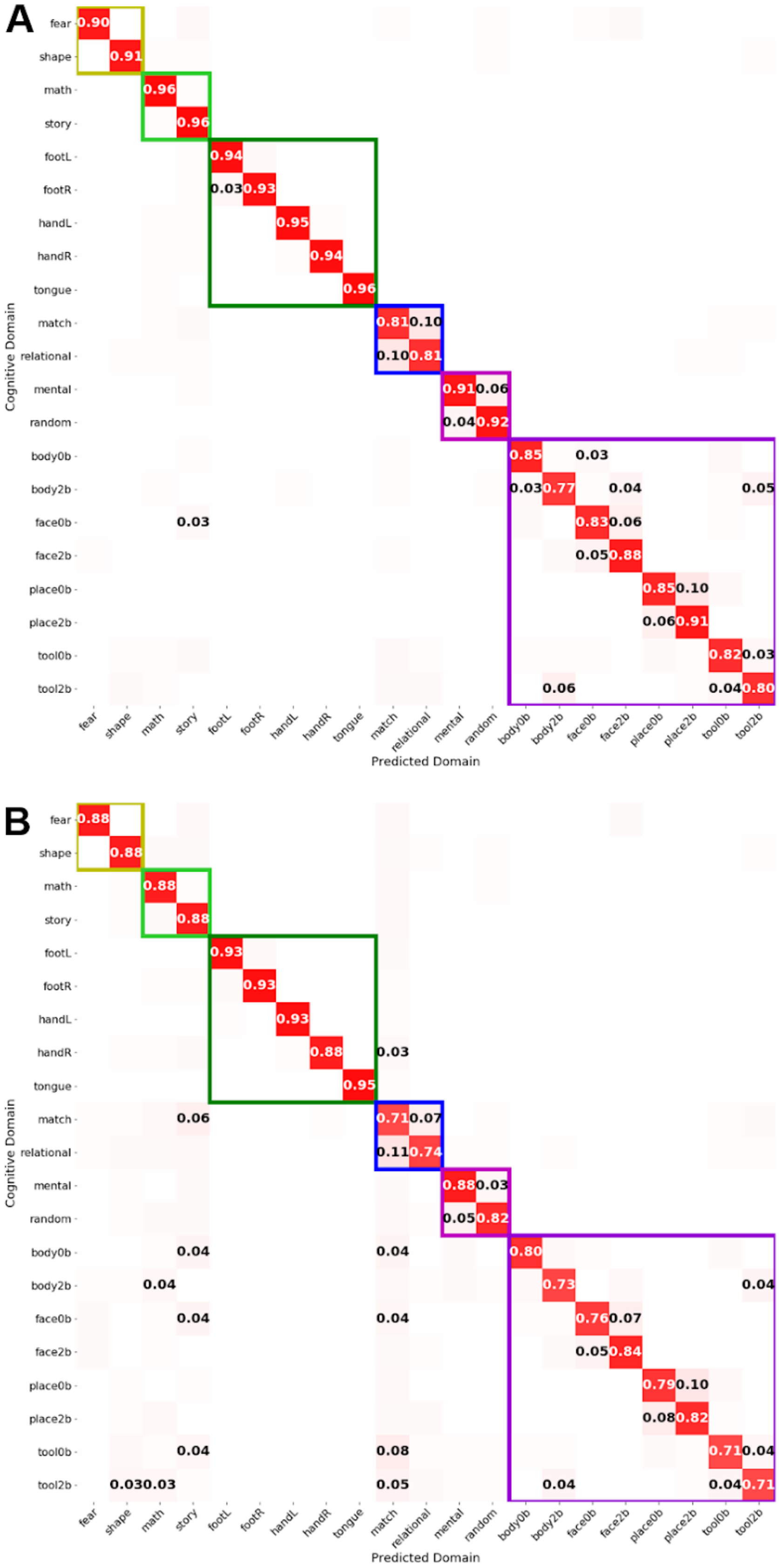


**Fig 2-Supplement 3. Confusion matrices of training a multidomain decoder (A) or combining multiple single-domain decoders (B).**

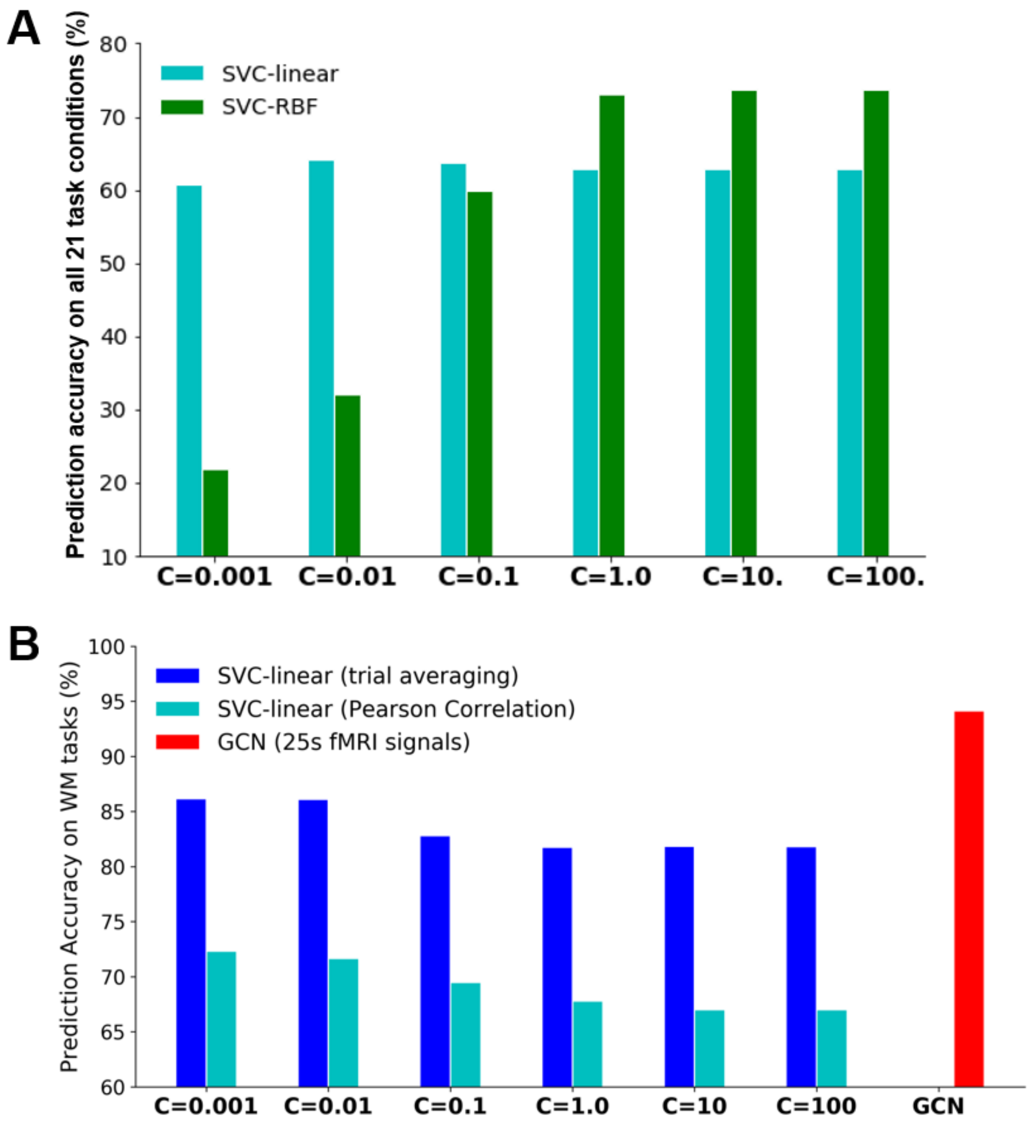


**Fig 2-Supplement 4. Comparison of decoding performance using different classifiers (A) or various types of features (B).**

First, on all task conditions (A), we tested different classifiers including SVC-linear and SVC-RBF with different regularization parameter C. A grid search approach was used to find the optimal C parameter (range: [0.001,0.01,0.1,1.,10.,100.]). The results indicated that the nonlinear kernel improved decoding accuracy for SVC (SVC-linear: 64.1%, SVC-RBF: 73.8%) while the linear model showed much lower capacity when decoding from fMRI activities.

Second, the impact of different features were investigated only on the WM task (B). Specifically, different features were tested for the SVC decoding models including averaging BOLD signals per task block (in yellow) and Pearson correlation of BOLD signals per task block (in cyan). Compared to these linear models, the GCN decoding model using short-series of fMRI signals (in red) achieved significantly better performance.

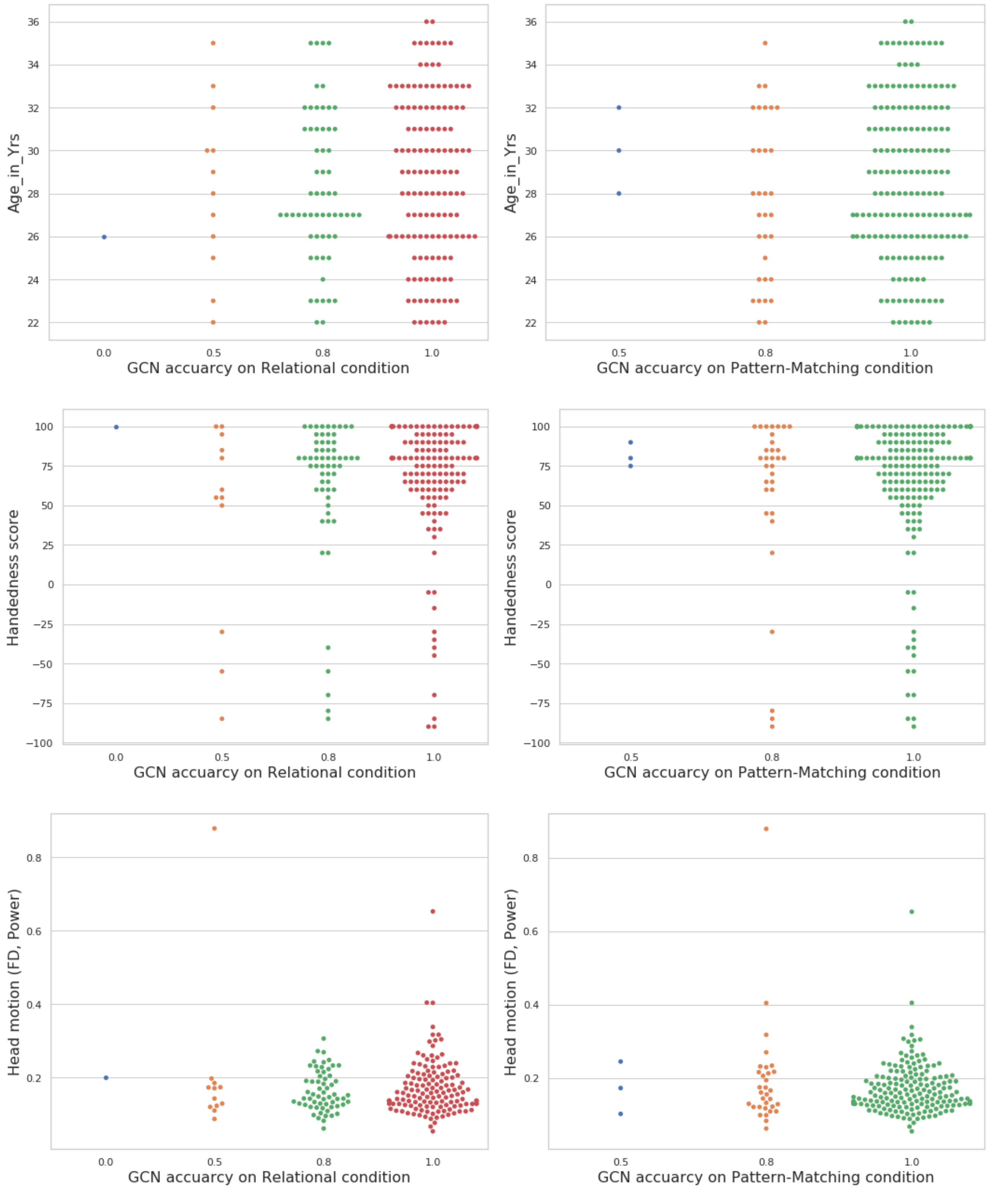


**Fig 3-Supplement 1. No association between subjects' age (first row), handedness (second row) and head motion (third row) and the GCN decoding accuracy on the relational processing task.**

We did not find significant association between GCN decoding accuracy on the relational processing tasks and subjects' age (r=0.04, p=0.50), handedness (r=0.02, p=0.75) and head motion (r=-0.07, p=0.29). The head motion was evaluated using framewise displacement (FD, Power) for each fMRI volume and averaged within each subject. To be noted that, the

F1-score was evaluated at the subject level for this analysis (2 functional runs, consisting of 6 relational blocks and 6 matching blocks). Due to such a small number of task trials per subject, the F1-score on each subject could only take a few specific values instead of a continuous distribution.

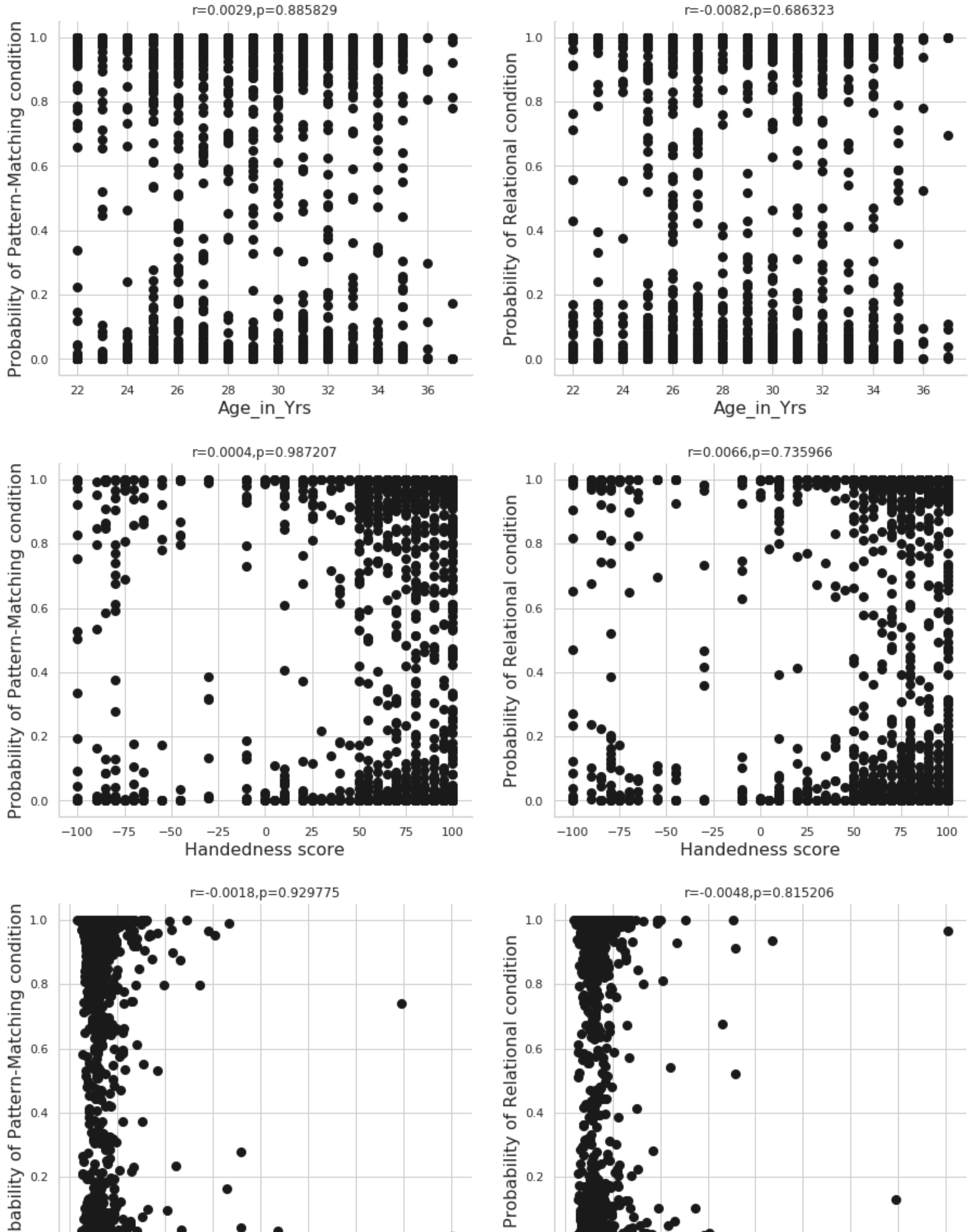


**Fig 3-Supplement 2. No association between subjects' age (first row), handedness (second row) and head motion (third row) and the model prediction (probability of each task condition) on the relational processing task, evaluated at the trial level.** We did not find a significant association between probability of each task condition (i.e. relational processing and pattern matching) and confounders at task trial level, including subjects' age (r=0.0029 p=0.88), handedness (r=0.0004, p=0.98) and head motion (r=-0.0018, p=0.92). To be noted that, since the age and handedness information were recorded at the subject level, we repeated the values for multiple times within each subject before correlation analysis. The head motion parameters were evaluated using framewise displacement (FD, Power) for each fMRI volume and then averaged within each task trial for the correlation analysis.

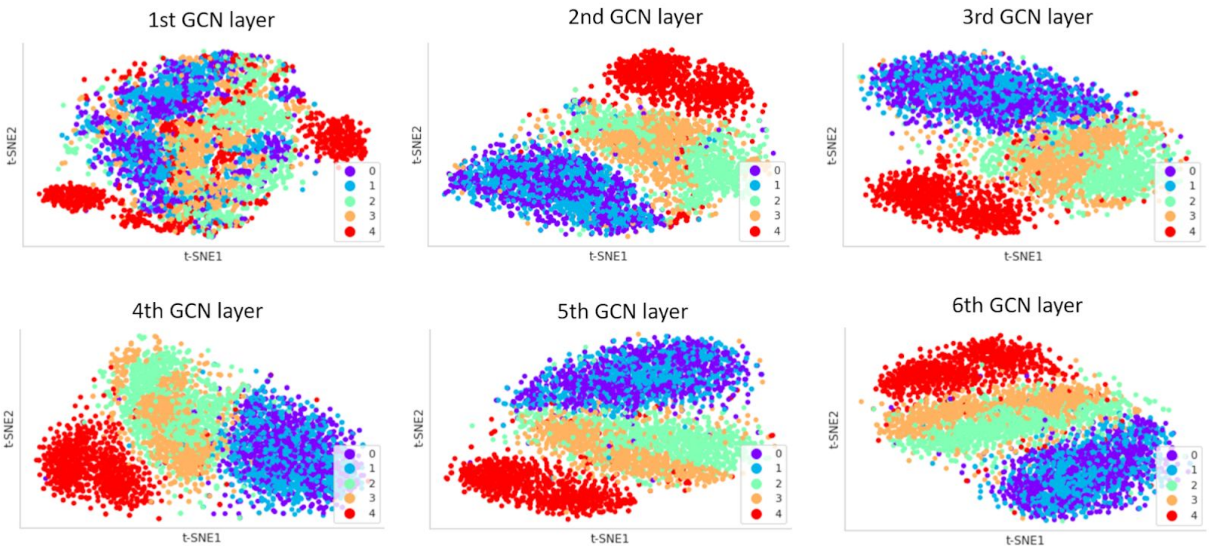


**Fig 4-Supplement 3. Visualization of layer activations for each GCN layer by projecting the learned representations onto 2-dimensional space by using t-SNE.**

The features of the five movement categories were randomly distributed in the raw fMRI data. No clear category structure was presented in the first GCN layer, highly mixed between different categories. In the 2nd and 3rd GCN layers, the tongue movement was easily distinguished from other tasks but still showing a high mixture effect between left and right movements. Finally, in the 5th and 6th GCN layers, all five categories of body movements were highly clustered and easily separated from each other.

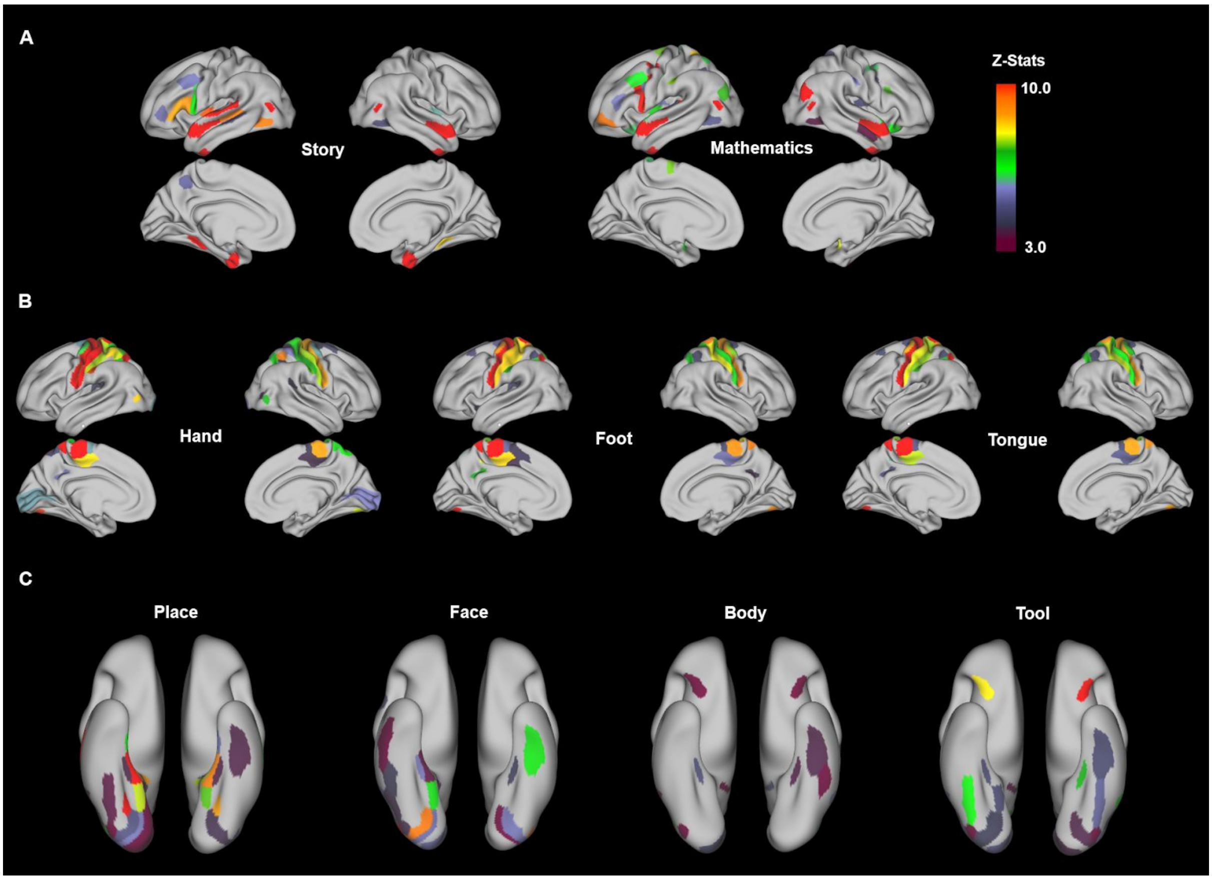


**Fig 5-Supplement 1. Meta-analysis of language, motor, and working memory tasks.**

Meta-analysis was conducted by searching for keywords in neuroquery (Dockès et al., 2020). For language task (A), we used the keyword “story” for language condition and “addition+subtraction” for the mathematical condition. For motor task (B), we used the keyword “hand movement” for hand condition, “foot+motor” for foot condition, “tongue+motor” for tongue condition. For the 0-back working memory task (C), we used the keyword “face recognition” for face condition, “body image” for body condition, “place+image” for place condition, “tool+image” for tool condition. The downloaded brain maps were first projected to the template surface “HCP_S1200_GroupAvg_v1 " using the ciftify tool (https://github.com/edickie/ciftify) and then mapped onto Glasser’s atlas (Glasser et al., 2016) for visualization. Only brain parcels with z-score above 3.0 were shown here to represent significant involvement of brain regions under the corresponding condition. Note that, the activation maps of the three conditions of the motor task were not easily differentiated here mainly due to the primary motor and somatosensory cortex being parcellated into single strips in the Glasser’s atlas (Glasser et al., 2016).

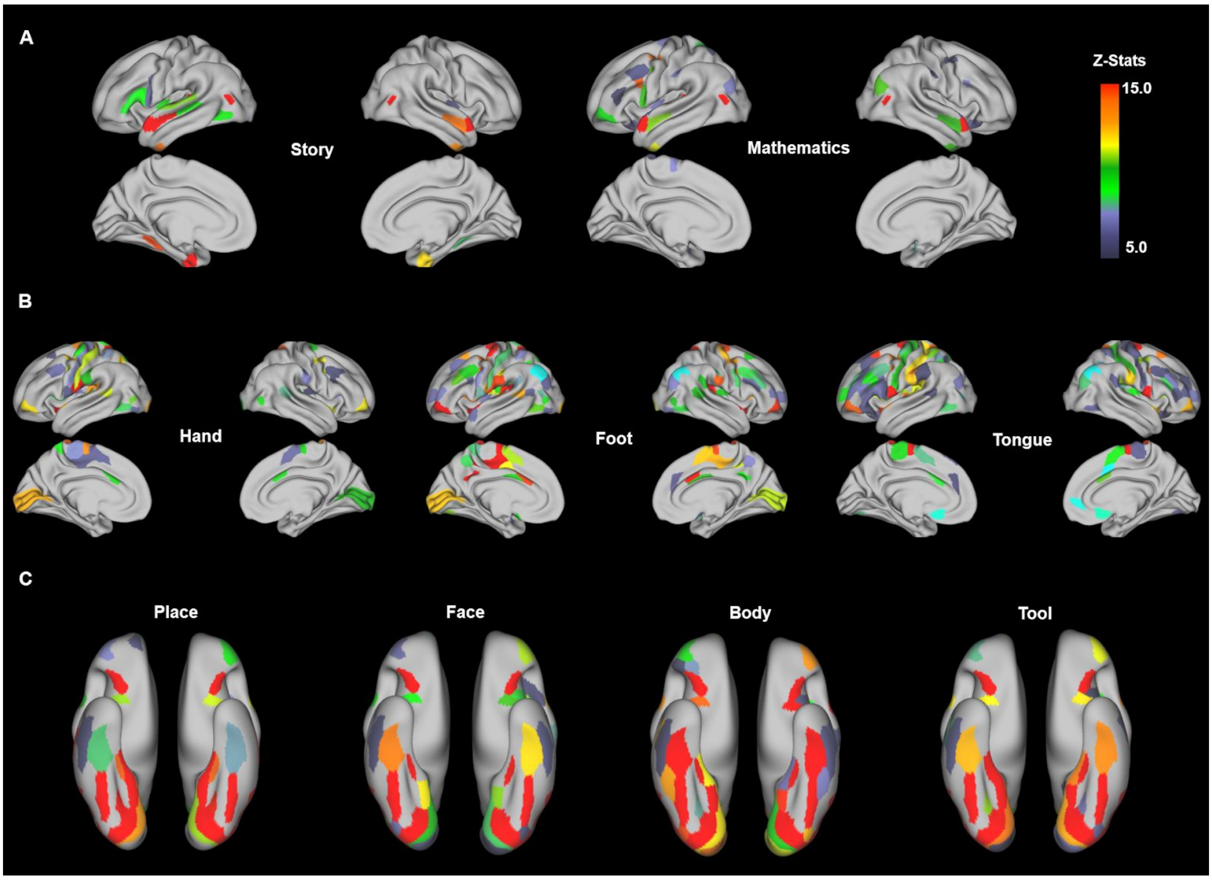


**Fig 5-Supplement 2. Activation maps of language, motor and working memory tasks from HCP.**

The contrast maps of HCP tasks (Barch et al., 2013) were downloaded from neurovault (https://neurovault.org/collections/457/), which contained a list of group-level z-stat maps for the task conditions. For language task (A), we showed the contrast of “Story vs Baseline” and “Math vs Baseline”. For motor task (B), we showed the contrast of “Right Hand vs Baseline”, “Right foot vs Baseline” and “Tongue vs Baseline”. For the 0-back working memory task (C), we showed the contrast of “0back Place vs Baseline”, “0back Face vs Baseline”, “0back Body vs Baseline” and “0back Tool vs Baseline”. The downloaded contrast maps were first projected to the template surface “HCP_S1200_GroupAvg_v1 " using the ciftify tool (https://github.com/edickie/ciftify) and then mapped onto Glasser’s atlas (Glasser et al., 2016) for visualization. Only brain parcels with z-score above 5.0 were shown here to represent strong brain activations under the corresponding condition.

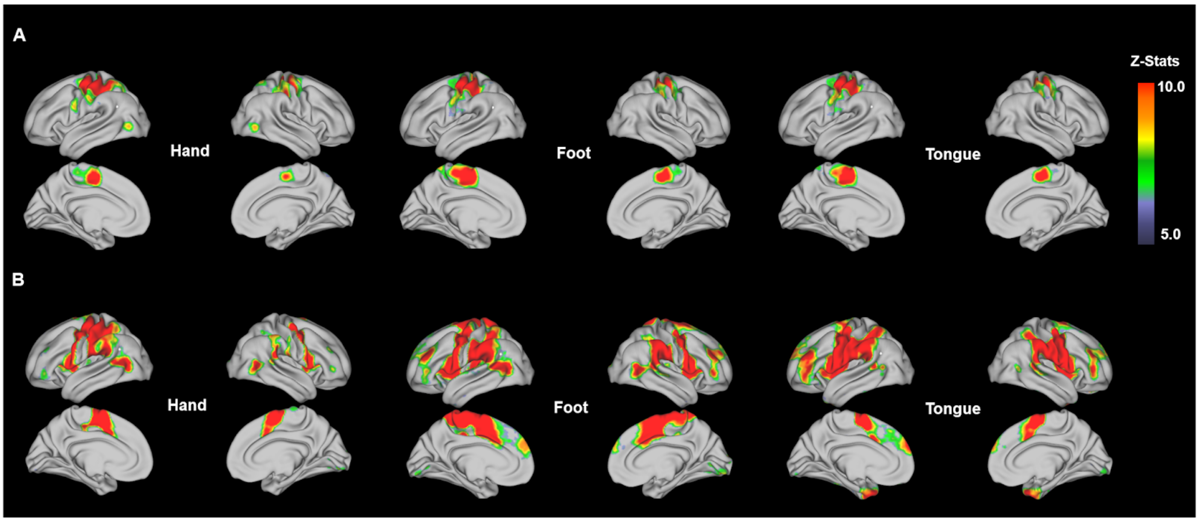


**Fig 5-Supplement 3. Meta-analysis and contrast maps for the motor task**

Meta-analysis (A) was conducted by searching for the keywords in neuroquery (Dockès et al., 2020). We used the keyword “hand movement” for hand condition, “foot+motor” for foot condition, “tongue+motor” for tongue condition. The contrast maps of HCP tasks (B) were downloaded from neurovault (https://neurovault.org/collections/457/). We only showed the contrast of “Right Hand vs Baseline”, “Right foot vs Baseline” and “Tongue vs Baseline” here. Both activation maps from meta-analysis and contrast maps from the HCP database were projected to the template surface “HCP_S1200_GroupAvg_v1 " for visualization.

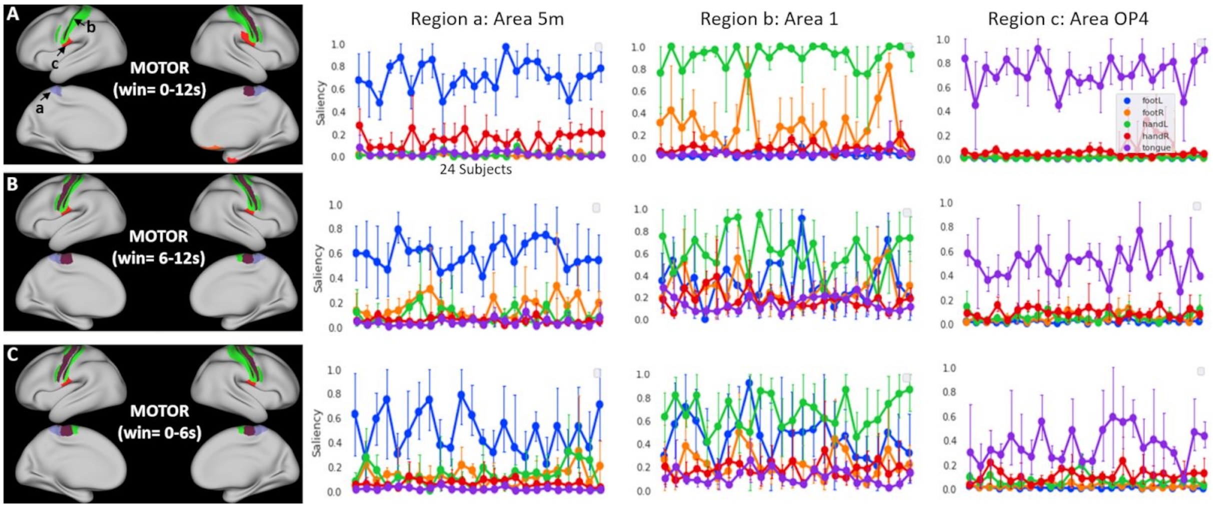


**Fig 5-Supplement 4. Stability of saliency maps among 480 motor trials extracted from 24 individual subjects.**

The stability of saliency maps were evaluated among different trials, between different subjects, and at different temporal resolutions. Specifically, we generated the saliency maps of the motor task using 24 subjects from the test set, including 2 functional runs for each subject, 10 motor trials within each run, in total of 480 individual task trials and consequently 480 saliency maps (one saliency map per trial). Three different models were evaluated using BOLD signals at different time windows, e.g. 0-12s, 6-12s and 0-6s (decoding accuracy: 97.33%, 95.53% and 86.54%). The results indicated highly consistent and category-specific patterns in the saliency maps. For instance, region a (labelled as “area 5m” in the Glasser atlas) selectively activated during foot movements, region b (area 1) selectively activated during hand movements, regions c (area OP4) selectively activated during tongue movements. When the window-size was shortened to 6-12s, although the model achieved a similar decoding accuracy (97.33% vs 95.53%), the inter-trial and inter-subject variability in saliency values in the four regions highly increased and might even interrupted the category-specific patterns among task conditions, especially for region b (area 1). Using the same window size but different time bins (i.e. 0-6s), the model achieved much lower decoding accuracy (86.54%). The saliency maps also indicated higher variability in all regions.

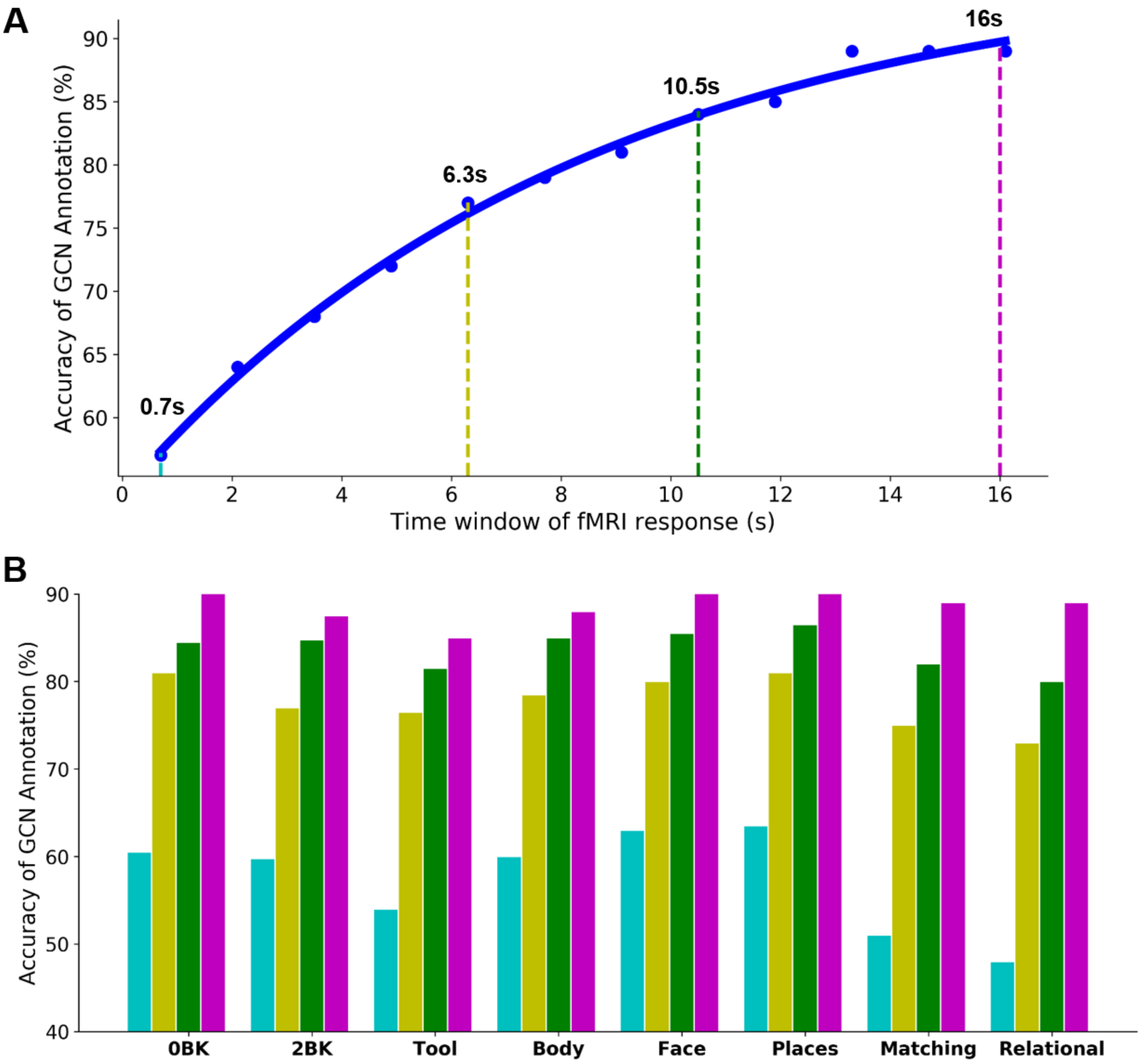


**Fig 6-Supplement 1. State annotation of relational processing and working memory conditions requires more than 10s to reach a plateau.**

The GCN model was trained based on the combination of all conditions from the relational processing and working memory tasks. With the minimal duration of working memory task trials lasting for 25s and relational reprocessing trials lasting for 16s, we evaluated the model with variable time windows, including a single fMRI volume (cyan: 0.7s), 9 TRs (yellow: 6.3s), 15 TRs (green: 10.5s) and 22 TRs (purple: 16s). Among all the experimental conditions, relational processing and recognition of tool images showed the lowest prediction scores at all levels of time windows.

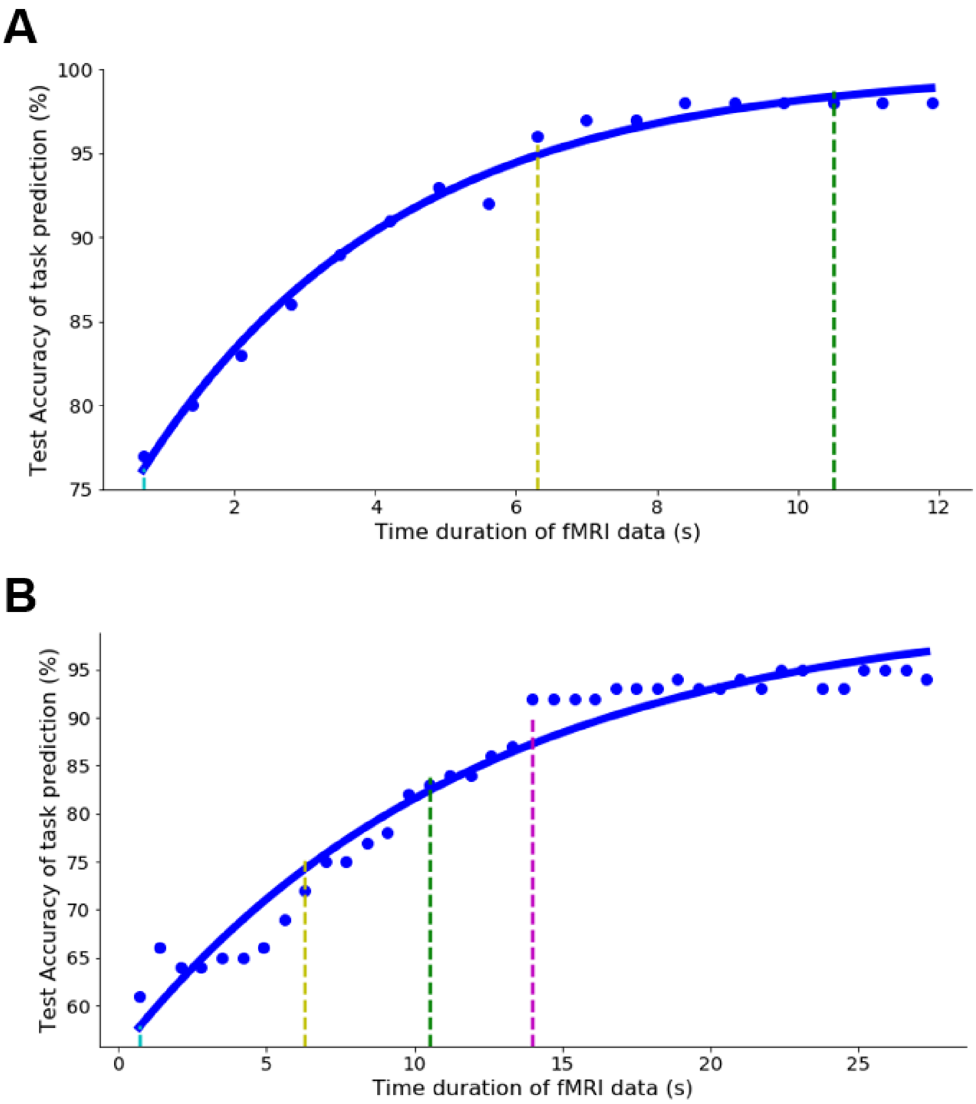


**Fig 6-Supplement 2. Decoding performance with variable temporal duration for Motor (A) and Working-memory tasks (B).**

For both task domains, the decoding accuracy gradually increased by using longer temporal duration and eventually plateaued at variable temporal durations. The point at which the model performance plateaued was not fixed but rather depending on the task design and the complexity of the target cognitive process. For instance, for Motor tasks, the decoding performance plateaued as early as 6s, while for Working-memory tasks, the performance plateaued at 14s.

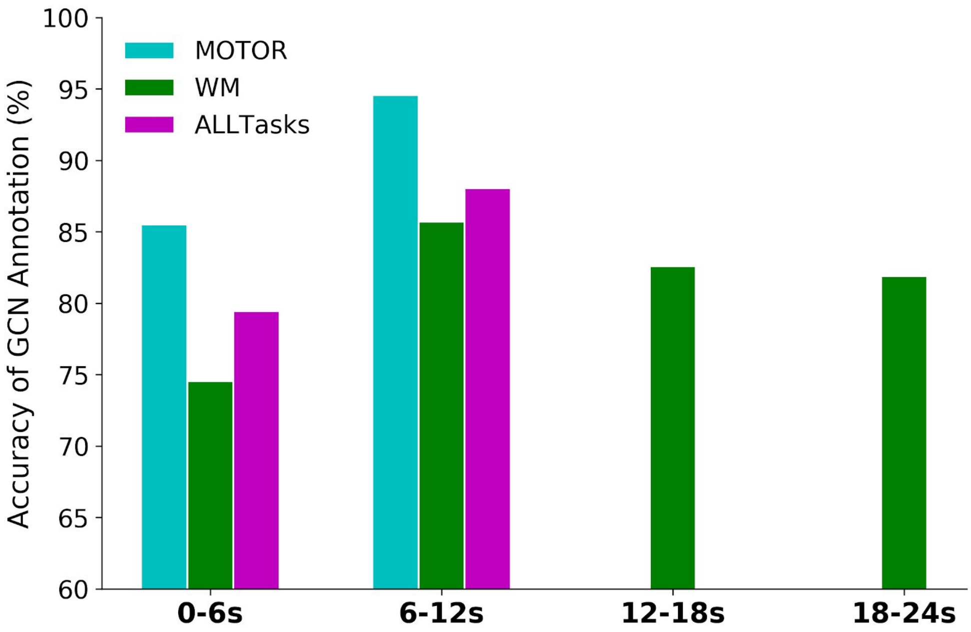


**Fig 7-Supplement 1. Performance of GCN annotation using a 6s window of fMRI signals.**

Task trials were split into mini-blocks with a temporal duration of 6s. Event blocks from the motor task last for 12s and thus were split into 2 mini-blocks of 6s time window. Event blocks from the working memory task last for 25s and thus were split into 4 mini-blocks of 6s time windows. These mini-blocks were treated as independent samples during model training. We also trained and evaluated separate decoding models for each of the time windows, by exclusively using the fMRI time series from the corresponding time bins.

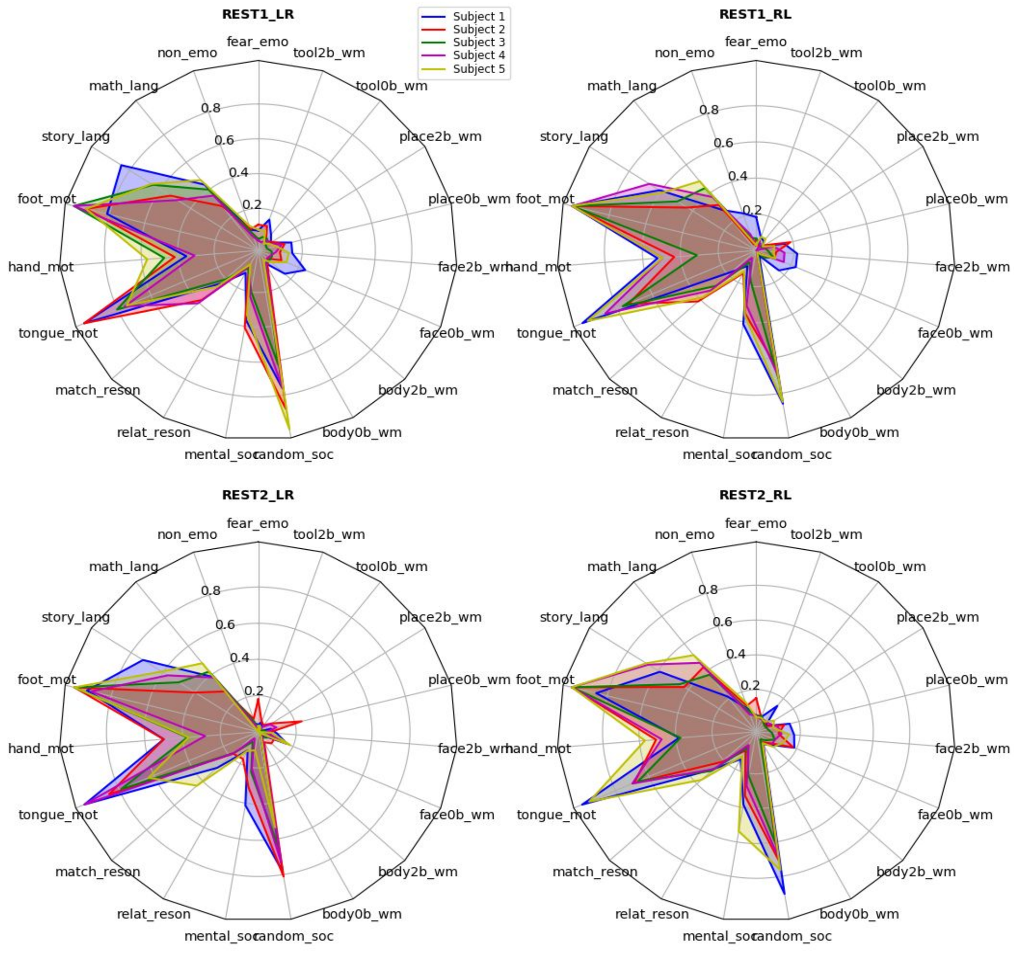


**Fig 7-Supplement 2. Distribution of dwelling time among the 21 predicted states using the resting-state fMRI dataset.**

We applied the pre-trained decoding model on 21 task conditions to predict the cognitive states using the resting-state fMRI time-series and plotted the proportion of dwell time for each cognitive state. The distribution of the 21 states was actually informative, for instance, the movement of foot and tongue, as well as non-social interaction (random_soc: watching clips of randomly moving shapes) were identified as the most frequent states. This result was somewhat expected because 1) body movement was unavoidable even in absence of cognitive process; 2) the random state somewhat represents mind-wandering activity during resting-state. However, these results should be interpreted with caution.

**Table S1: Comparison of decoding performance between different models.**
We reported the best performance for the baseline models after a grid search of the hyperparameters. For GCN-avg-trial (using trial-averaged BOLD signal as model inputs) and GCN (using fMRI time-series as model inputs), we reported the mean and standard deviation of the decoding accuracies among 10 fold cross-validation with shuffle splits.

| **Models** | **Train Accuracy** | **Validation Accuracy** | **Test Accuracy** |
|---|---|---|---|
| SVC-linear | 67.2% | 63.3% | 64.1% |
| SVC-rbf | 99.7% | 73.5% | 73.8% |
| Random Forest | 100% | 48.0% | 47.5% |
| GCN-avg-trial | 86.9%(+/-0.39%) | 77.7%(+/-0.2%) | 78.8%(+/-0.21%) |
| GCN | 96.3%(+/-0.42%) | 90.2%(+/-0.21%) | 90.7%(+/-0.20%) |

**SI Appendix: Legends for supporting videos**

**Fig 4-Supplement 1. Projection of raw fMRI data and learned representations from the decoding model.**

To illustrate the effect of graph convolution, we projected both raw fMRI time-series and learned representations from graph convolutional networks onto a 2-dimensional space by using different dimension reduction techniques, including PCA (first column), t-SNE (second column) (Maaten and Hinton, 2008), UMAP (third column) (McInnes et al., 2018), and PHATE (last column) (Moon et al., 2019). The data samples includes five types of movements, i.e. the movement of right foot (class 0, in red), left foot (class 1, in cyan), right hand (class 2, in green), left hand (class 3, in blue), and tongue (class 4, in purple). The five types of movements were not separable and highly overlapped in either fMRI time-series or during the early stages of training process (before Epoch #10) by using either of the four projection methods. A clustering effect started to show after 20 epochs of training, that samples of the same type of movements were located closest to each other while well-separated from other classes. Best visualization was provided by t-SNE.

**Fig 4-Supplement 2. Visualization of learned representations of Motor task-fMRI data during the training process.**

The layer activations of the last graph convolutional layer of the decoding model were extracted as the graph representations of brain dynamics. The learned representations were then projected onto a 2-dimensional space using t-SNE (left panel) (Maaten and Hinton, 2008). The data samples includes five types of movements, i.e. the movement of right foot (class 0, in red), left foot (class 1, in cyan), right hand (class 2, in green), left hand (class 3, in blue), and tongue (class 4, in purple). The decoding accuracies on the training (blue line) and test (red line) sets were also collected during the model training process and shown in the right panel. The chance level of classifying the five types of movements was 20%, marked as a cyan line in the plot. Note that, the decoding performance started at the chance level of 20% and quickly reached the plateau of 97% after 20 training epochs.

### References

Barch, D.M., Burgess, G.C., Harms, M.P., Petersen, S.E., Schlaggar, B.L., Corbetta, M., Glasser, M.F., Curtiss, S., Dixit, S., Feldt, C., Nolan, D., Bryant, E., Hartley, T., Footer, O., Bjork, J.M., Poldrack, R., Smith, S., Johansen-Berg, H., Snyder, A.Z., Van Essen, D.C., WU-Minn HCP Consortium, 2013. Function in the human connectome: task-fMRI and individual differences in behavior. Neuroimage 80, 169–189.

Dockès, J., Poldrack, R.A., Primet, R., Gözükan, H., Yarkoni, T., Suchanek, F., Thirion, B., Varoquaux, G., 2020. NeuroQuery, comprehensive meta-analysis of human brain mapping. Elife 9. https://doi.org/10.7554/eLife.53385

Glasser, M.F., Coalson, T.S., Robinson, E.C., Hacker, C.D., Harwell, J., Yacoub, E., Ugurbil, K., Andersson, J., Beckmann, C.F., Jenkinson, M., Smith, S.M., Van Essen, D.C., 2016. A multi-modal parcellation of human cerebral cortex. Nature 536, 171–178.

Maaten, L. van der, Hinton, G., 2008. Visualizing Data using t-SNE. J. Mach. Learn. Res. 9, 2579–2605.

McInnes, L., Healy, J., Saul, N., Großberger, L., 2018. UMAP: Uniform Manifold Approximation and Projection. Journal of Open Source Software. https://doi.org/10.21105/joss.00861

Moon, K.R., van Dijk, D., Wang, Z., Gigante, S., Burkhardt, D.B., Chen, W.S., Yim, K., van den Elzen, A., Hirn, M.J., Coifman, R.R., Ivanova, N.B., Wolf, G., Krishnaswamy, S., 2019. Visualizing structure and transitions in high-dimensional biological data. Nat. Biotechnol. 37, 1482–1492.
